# MIF-2/D-DT is an atypical atherogenic chemokine that promotes advanced atherosclerosis and hepatic lipogenesis

**DOI:** 10.1101/2021.12.28.474328

**Authors:** Omar El Bounkari, Chunfang Zan, Jonas Wagner, Elina Bugar, Priscila Bourilhon, Christos Kontos, Marlies Zarwel, Dzmitry Sinitski, Jelena Milic, Yvonne Jansen, Wolfgang E. Kempf, Lars Mägdefessel, Adrian Hoffmann, Markus Brandhofer, Richard Bucala, Remco T. A. Megens, Christian Weber, Aphrodite Kapurniotu, Jürgen Bernhagen

## Abstract

Atherosclerosis is the underlying cause of cardiovascular diseases (CVDs) such as myocardial infarction and ischemic stroke. It is a lipid-triggered chronic inflammatory condition of the arterial vascular wall that is driven by various inflammatory pathways including atherogenic cytokines and chemokines. D-dopachrome tautomerase (D-DT), also known as macrophage migration inhibitory factor-2 (MIF-2), belongs to the MIF protein family, which is best known for its pathogenic role in a variety of inflammatory and immune conditions including CVDs. While MIF is well known as a promoter of atherogenic processes, MIF-2 has not been studied in atherosclerosis. Here, we investigated atherosclerosis in hyperlipidemic *Mif-2^−/−^Apoe^−/−^* mice and studied the role of MIF-2 in various atherogenic assays *in vitro*. We found that global *Mif-2* deficiency as well as its pharmacological blockade by 4-CPPC protected against atherosclerotic lesion formation and vascular inflammation in models of early and advanced atherogenesis. On cellular level, MIF-2 promoted monocyte migration in 2D and 3D and monocyte arrest on aortic endothelial monolayers, promoted B-cell chemotaxis *in vitro* and B-cell homing *in vivo*, and increased macrophage foam cell formation. Dose curves and direct comparison in a 3D migration set-up suggest that MIF-2 may be a more potent chemokine than MIF for monocytes and B cells. We identify CXCR4 as a novel receptor for MIF-2. The evidence relies on a CXCR4 inhibitor, CXCR4 internalization experiments, MIF-2/CXCR4 binding studies by yeast-CXCR4 transformants, and fluorescence spectroscopic titrations with a soluble CXCR4 surrogate. Of note, *Mif-2^−/−^Apoe^−/−^* mice exhibited decreased plasma cholesterol and triglyceride levels, lower body weights, smaller livers, and profoundly reduced hepatic lipid accumulation compared to *Apoe^−/−^* mice. Mechanistic experiments in Huh-7 hepatocytes suggest that MIF-2 regulates the expression and activation of sterol-regulatory element binding protein-1 and −2 (SREBP-1, SREBP-2) to induce lipogenic downstream genes such as FASN and LDLR, while it attenuated the activation of the SREBP inhibiting AMPK pathway. Studies using receptor Inhibitors showed that SREBP activation and hepatic lipoprotein uptake by MIF-2 is mediated by both CXCR4 and CD74. Lastly and in line with a combined role of MIF-2 in vascular inflammation and hepatic lipid accumulation, MIF-2 was found to be profoundly upregulated in unstable human carotid plaques, underscoring a critical role for MIF-2 in advanced stages of atherosclerosis. Together, these data identify MIF-2 as a novel atherogenic chemokine and CXCR4 ligand that not only promotes lesion formation and vascular inflammation but also strongly affects hepatic lipogenesis in an SREBP-mediated manner, possibly linking atherosclerosis and hepatic steatosis.

## INTRODUCTION

Cardiovascular diseases (CVDs) including myocardial infarction and ischemic stroke remain the leading cause of mortality in Western societies and are responsible for an estimated 30% of all deaths worldwide. Atherosclerosis is the main underlying cause of CVDs and a lipid-triggered chronic inflammatory condition of the arterial vascular wall that is driven by various immune and inflammatory pathways (1–4).

Atherosclerotic diseases are promoted by risk and lifestyle factors such as hypertension, smoking, high-fat Western-type diets, and lack of physical activity. Also, disease pathology has been associated with co-morbidities such type 2 diabetes (T2D) or non-alcoholic fatty liver disease (NAFLD) (5). Triggered by oxidized low density lipoprotein (oxLDL) and arterial endothelial stress, atheroprogression involves leukocyte infiltration and lesional inflammation, processes that are chiefly orchestrated and promoted by the dysregulated activity of cytokines and chemokines, acting in a mostly sequential but also pleiotropic and redundant manner (4, 6). For example, classical chemokines such as CCL2 or CXCL1/CXCL8 promote the atherogenic arrest, transmigration, and intimal accumulation of monocytes via CCR2 and CXCR2, and amplify lesional inflammation by a variety of mechanisms.

Macrophage migration inhibitory factor (MIF) is an evolutionarily conserved multi-functional inflammatory mediator with important pathogenic roles in acute and chronic inflammatory diseases (7–10). Plasma MIF levels are associated with coronary artery disease (CAD) and we previously detected abundant MIF expression in atherosclerotic plaques (11–14) and demonstrated that MIF promotes the atherogenic recruitment of monocytes and T cells through non-cognate interaction with the CXC chemokine receptors CXCR2 and CXCR4 (15). The MIF/CXCR4 axis may be of particular relevance in atherosclerosis, as it has been associated with atherogenic activities of not only monocytes and T cells, but also neutrophils, platelets, and B cells (15–18). Accordingly, MIF has been designated an atypical chemokine (ACK), an emerging family of structurally diverse proteins with chemokine-like properties that bind to classical chemokine receptors with high-affinity, while lacking classifying structural features of chemokines such as N-terminal cysteines and a chemokine-fold (9, 19). MIF also binds to CD74, the cognate receptor of MIF and a type II transmembrane glycoprotein, additionally known as the plasma membrane form of MHC class II invariant chain Ii (20). MIF engagement of CD74 or CXCR4 can occur by direct high-affinity binding (15, 20), but depending on the cellular context, MIF signaling through CD74 requires the co-receptor CD44 (21, 22) and biochemical evidence suggests the formation of receptor complexes between CD74 and MIF’s chemokine receptors as a molecular basis for altered signaling responses that can serve to augment the migration of lymphocytes and myeloid cells (15, 16, 23, 24). Thus, while MIF’s pro-atherogenic activities are mainly mediated through its chemokine receptor pathways, CD74 directly and indirectly contributes to MIF’s role in vascular inflammation. However, the MIF/CD74 axis may also lead to the activation of adenosine-monophosphate kinase (AMPK), a pathway that has been associated with MIF-mediated tissue protection in cardiac ischemia and hepatosteatosis (25, 26).

D-dopachrome tautomerase (D-DT) is a member of the MIF protein family and was originally identified as a cytoplasmic enzyme in human melanoma cells and liver tissue that can convert the non-natural substrate D-dopachrome to 5,6-dihydroxyindole (27, 28). D-DT consists of 118 amino acids and shares 34% sequence homology and a remarkably similar three-dimensional structural architecture with MIF (29). Despite its identification over 25 years ago, a role for D-DT in human immunity only emerged fairly recently, when D-DT was characterized as a cytokine and functional homolog of MIF (30). Accordingly, D-DT is now also termed MIF-2 (30–32). MIF-2 is expressed in numerous organs such as liver, heart, lung, and pancreas. Compared to MIF, the mechanism of action of MIF-2 is incompletely understood. MIF-2 interacts with CD74 with high affinity to promote activation of the MAPK pathway, thus sharing with MIF this cognate receptor signaling pathway (30). Moreover, recent evidence implicates MIF-2/ACKR3 interactions in lung epithelial repair in chronic obstructive pulmonary disease (COPD), but whether this effect is based on direct or indirect binding requires further scrutiny (33–35). MIF-2 lacks the pseudo(E)LR motif required for interaction between MIF and its non-cognate receptor CXCR2 (15, 36), and accordingly does not interact with CXCR2 (37). Furthermore, functional studies in disease models suggest that, depending on the cell type, tissue, or disease context, MIF and MIF-2 may exhibit overlapping or different regulatory effects. For example, MIF-2 promotes renal cell carcinoma and endotoxemia in a similar manner as MIF (30, 38, 39), while MIF but not MIF-2 recruits inflammatory macrophages in an experimental polymicrobial sepsis model (37). Interestingly, in adipogenesis and adipose tissue inflammation, MIF and MIF-2 have been reported to display opposite activities (40, 41). In the same vein, MIF levels positively correlate with obesity outcomes, whereas MIF-2 levels show a negative correlative relationship (42, 43). In ischemic heart disease and heart failure, both MIF and MIF-2 have been found to activate the AMPK pathway via the CD74/CD44 axis to exert cardioprotective effects (26, 44–47), However, increased MIF levels during cardiac surgery were associated with organ-protective properties during myocardial I/R, while high MIF-2 levels were predictive of the development of organ dysfunction (48), altogether suggesting a complex role of MIF proteins in heart disease. In contrast to MIF, the role of MIF-2 in the chronic pathophysiological process of atherosclerosis has been not explored so far.

Applying a model of hyperlipidemic atherogenic *Apolipoprotein E*-deficient (*Apoe^−/−^*) mice in combination with global *Mif-2* deficiency or MIF-2 selective pharmacological blockade, here we wished to investigate the role of MIF-2 in atherosclerosis *in vivo*. *Mif-2^−/−^Apoe^−/−^* mice on high-fat diet (HFD) exhibited reduced atherosclerotic lesions in models of both early and advanced atherogenesis compared to Mif-2-proficient *Apoe^−/−^* control mice. *In vitro* flow arrest, chemotaxis, and *in vivo* recruitment experiments indicated that Mif-2 behaved as a chemokine for monocytes and B cells, and, compared to MIF, was the more potent chemokine by high affinity-engagement of CXCR4. Strikingly, atherogenic *Mif-2^−/−^Apoe^−/−^* mice showed reduced plasma levels of cholesterol, triglycerides, and atherogenic lipoproteins, and were protected from hepatosteatosis. Sterol regulatory element binding proteins (SREBPs) are endoplasmic reticulum (ER) membrane-bound transcription factors that are activated by proteolytic processing and nuclear translocation and play a central role in controlling the expression of several genes involved in hepatic lipogenesis and lipid homeostasis (49, 50). Mechanistic *in vitro* experiments employing the human hepatocyte cell line Huh-7, SREBP and SREBP target gene analysis, receptor blocking agents, proximity ligation assay (PLA), and FLIM/FRET analysis suggested that MIF-2 acts as a lipogenic factor that promotes hepatic lipogenesis through activation of the CXCR4/CD74-SREBP axis. In conjunction with an identified prominent upregulation of MIF-2 in unstable human atherosclerotic plaques from carotid endarterectomies (CEAs), our data suuport a role for MIF-2 in advanced atherosclerosis that is furthered by a dual lipid/hepatic and vascular phenotype.

## RESULTS

### Genetic deletion and pharmacological blockade of MIF-2 mitigate atherosclerotic lesion formation in early stages of atherosclerosis

To explore the potential involvement of MIF-2 in the pathogenesis of atherosclerosis, we first assessed the impact of its genetic deletion on disease progression in a mouse model of early atherogenesis. *Mif-2^−/−^Apoe^−/−^* mice were generated by cross-breeding *Mif-2^+/–^* and athero-genesis-prone *Apoe^−/−^* mice (Figure S1A-D). Groups of female *Mif-2^−/−^Apoe^−/−^ versus Apoe^−/−^* littermates (8-week-old) were subsequently fed a Western-type cholesterol-rich high-fat diet (HFD) for 4.5 weeks to initiate atherosclerotic lesion formation (Figure 1A). *Mif-2-*deficient *Apoe^−/−^* mice displayed no gross phenotype differences compared to their *Apoe^−/−^* littermates. Interestingly, comparison of blood cell counts revealed that *Mif-2^−/−^Apoe^−/−^* mice exhibited a decreased monocyte and slightly increased blood T-cell count (Table S1).

**Figure 1.**
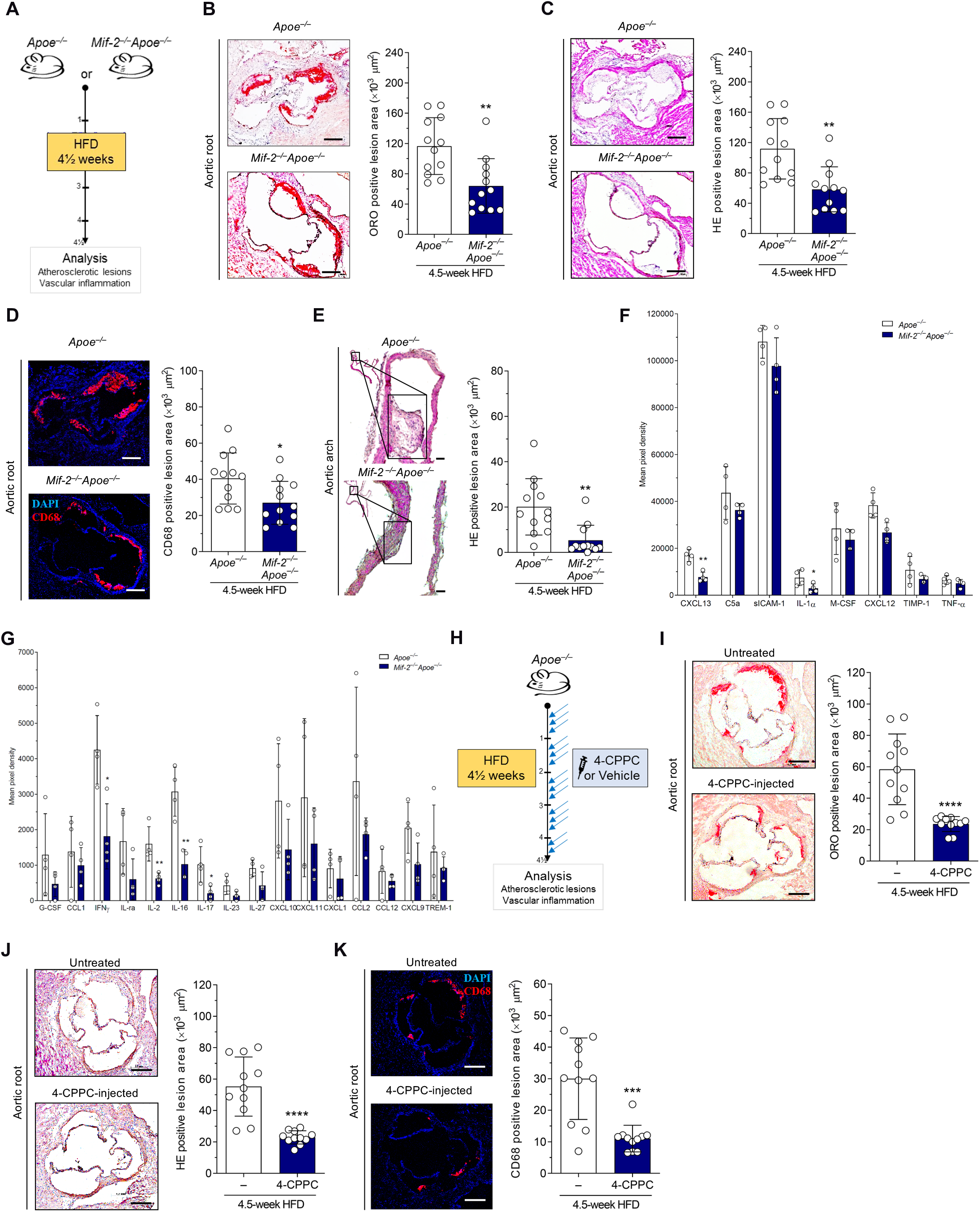
Genetic deletion or pharmacological inhibition of MIF-2 attenuates early atherosclerotic lesion progression and vascular inflammation *in vivo.* (A) Experimental outline. Female *Apoe^−/−^* and *Mif-2^−/−^Apoe^−/−^* mice were fed a high-fat diet (HFD) containing 0.21% cholesterol for 4.5 weeks to develop early-stage plaques. (B) Representative Oil Red O (ORO) staining images of aortic roots in frozen sections (8 µm) from *Apoe^−/−^* and *Mif-2^−/−^ Apoe^−/−^* mice and corresponding quantification results (12 serial sections per mouse). n = 12 mice; each data point represents one mouse; scale bar, 250 µm. (C) Hematoxylin-eosin (HE) staining images of aortic roots in frozen sections (8 µm) from *Apoe^−/−^* and *Mif-2^−/−^Apoe^−/−^* mice and corresponding quantification results (12 serial sections per mouse). n = 12 mice; each data point represents one mouse; scale bar, 250 µm. (D) Representative images and quantification results of CD68^+^ macrophages (red) in aortic root sections (8 µm) from *Apoe^−/−^* and *Mif-2^−/−^Apoe^−/−^* mice (DAPI, blue). n = 12 mice; each data point represents one mouse; scale bar, 250 µm. (E) Representative HE-stained images of aortic arch in paraffin sections (4 µm) and quantification results (8 sections per mouse). n = 12 mice; each data point represents one mouse; scale bar, 250 µm. (F, G) Quantitative analysis by mouse cytokine array featuring 40 inflammatory/atherogenic cytokines/chemokines on plasma samples from both groups of mice. Data are means ± SD from 4 mice per group, performed in duplicate each. (F) Cytokines/chemokines with a relatively higher plasma abundance; (G) cytokines/chemokines with a relatively lower plasma abundance. (H) Experimental outline: *Apoe^−/−^* mice were fed a HFD for 4.5 weeks and administrated 4-CPPC (50 µg/mouse; 3x per week) or vehicle (3x per week) in parallel. (I) Representative ORO staining images of aortic roots in frozen serial sections (8 µm) from *Apoe^−/−^* mice administrated with 4-CPPC or vehicle and corresponding quantification results (12 sections per mouse; n = 11 mice per group). Scale bar, 250 µm. (J) Same as in (I) except that HE staining was performed. (K) Representative images and quantification results of CD68^+^ macrophage content (red) in aortic root sections (8 µm) from *Apoe^−/−^* mice administrated with 4-CPPC or vehicle (DAPI, blue). n = 11 mice per group. Scale bar, 250 µm. All values are means ± SD; *p<0.05; **p<0.01; ***p<0.001; ****p<0.0001.

Quantification of atherosclerotic lesions following oil-red O (ORO) and hematoxylin-eosin (HE) staining revealed significantly reduced lesion areas in cross-sections of aortic root and vertical sections of aortic arch with an approximate reduction of 50% observed in the *Mif-2^−/−^Apoe^−/−^* mice compared to *Apoe^−/−^* controls (Figures 1B, C, and E). Moreover, reduced plaque formation was accompanied by a substantially decreased number of accumulated lesional macrophages in *Mif-2^−/−^Apoe^−/−^* mice compared to *Apoe^−/−^* controls as determined by lesional CD68 staining (Figure 1D), and in line with the observed decrease in circulating monocytes (Table S1). As vascular inflammation correlates with the expression of inflammatory mediators during atherogenesis, we evaluated the plasma levels of inflammatory cytokines by a protein array approach (Figures 1F, G, and S2). We determined a significant reduction of IFN-γ, IL-2, IL-16, IL-17, CXCL13, and IL-1α concentrations in *Mif-2^−/−^Apoe^−/−^* mice compared with *Apoe^−/−^* controls, while trends for reductions of several other cytokines were observed as well (Figure 1F-G). Of note, a similar atherogenic phenotype was also observed in male *Mif-2^−/−^Apoe^−/−^*mice on a 4.5-week HFD, with smaller atherosclerotic plaques in aortic root and decreased vascular inflammation when compared to *Apoe^−/−^* control mice (Figure S3A-D). Collectively, these results indicate that global *Mif-2* deficiency attenuates early atherogenesis and vascular inflammation in *Apoe^−/−^* mice of both sexes.

To further confirm the role of MIF-2 in early atherogenesis and address the potential role of compensatory effects caused by global gene deficiency, we employed a pharmacological blockade approach, using 4-CPPC, a specific small molecule inhibitor of MIF-2 that exhibits a 13-fold selectivity against MIF-2 over MIF (32, 51). *Apoe^−/−^* mice were placed on a 4.5-week HFD and 4-CPPC *versus* vehicle control was administered in parallel to the HFD (intraperitoneal (i.p.) administration; 2.5 mg kg^−1^) three times per week (Figure 1H). Corroborating the data obtained in the genetic deletion model, atherosclerotic lesion size in aortic root was markedly decreased by approximately 50% in 4-CPPC-treated mice compared with vehicle-treated controls as analyzed by ORO and HE staining (Figure 1I-J). Similarly, there was a pronounced reduction in lesional macrophages in the 4-CPPC-treated group (Figure 1K). Taken together, the genetic deletion and pharmacological blockade models suggest that MIF-2 is a novel pro-atherogenic player in early stages of atherosclerosis.

### MIF-2 promotes leukocyte adhesion and chemotactic migration

The contribution of MIF-2 to the progression of early stages of atherosclerosis raised the possibility that, similar to MIF, MIF-2 may also exert atherogenic properties of leukocytes by influencing leukocyte adhesion and migration. To address this possibility, we first assessed the adhesion of MonoMac6 monocytes to a confluent monolayer of human aortic endothelial cells (HAoECs) under flow stress conditions. Recombinant MIF-2 dose-dependently promoted the cell arrest of MonoMac6 cells on HAoECs, with a peak effect observed at a concentration of 1.6 nM and a significant upregulation was also noted at 0.8 and 4 nM MIF-2. Of note, the arrest effect of 1.6 nM MIF-2 was higher than that of 16 nM MIF, which was analyzed for comparison (Figure 2A), indicating that MIF-2 has a more potent arrest capacity as MIF. Next, we interrogated the migratory capacity of primary human monocytes in response to MIF-2 employing a Transwell-based chemotaxis set-up. As shown in Figure 2B, MIF-2 elicited the chemotactic migration of monocytes in a dose-dependent manner, exhibiting a bell-shaped dose curve typically seen for chemokines, with an optimum at 4 nM. This peak concentration is thus well in line with the concentration optimum of MIF-2 seen in the arrest assay. To further verify this effect, we examined the impact of MIF-2-selctive small molecule inhibitor 4-CPPC on monocyte migration triggered by the optimal MIF-2 dose under the same conditions. 4-CPPC potently and dose-dependently inhibited monocyte migration in response to MIF-2, with complete ablation of the MIF-2 effect already seen at a 5-fold molar excess of the inhibitor (Figure 2C). Additionally, we investigated MIF-2-mediated monocyte migration in a three-dimensional (3D) collagen-based model applying time-lapse microscopy and single cell tracking. To this end, primary human monocytes were seeded into collagen gels and subjected to different concentrations of a MIF-2 gradient. MIF-2 enhanced the motility and directional migration of monocytes in a concentration-dependent fashion as illustrated by the corresponding single-cell migration tracks (Figure 2D), and quantitative analysis of the forward migration index (Figure 2E). Together, these data therefore provided evidence that MIF-2 is a chemokine-like cytokine that promotes monocyte adhesion and chemotactic migration.

**Figure 2.**
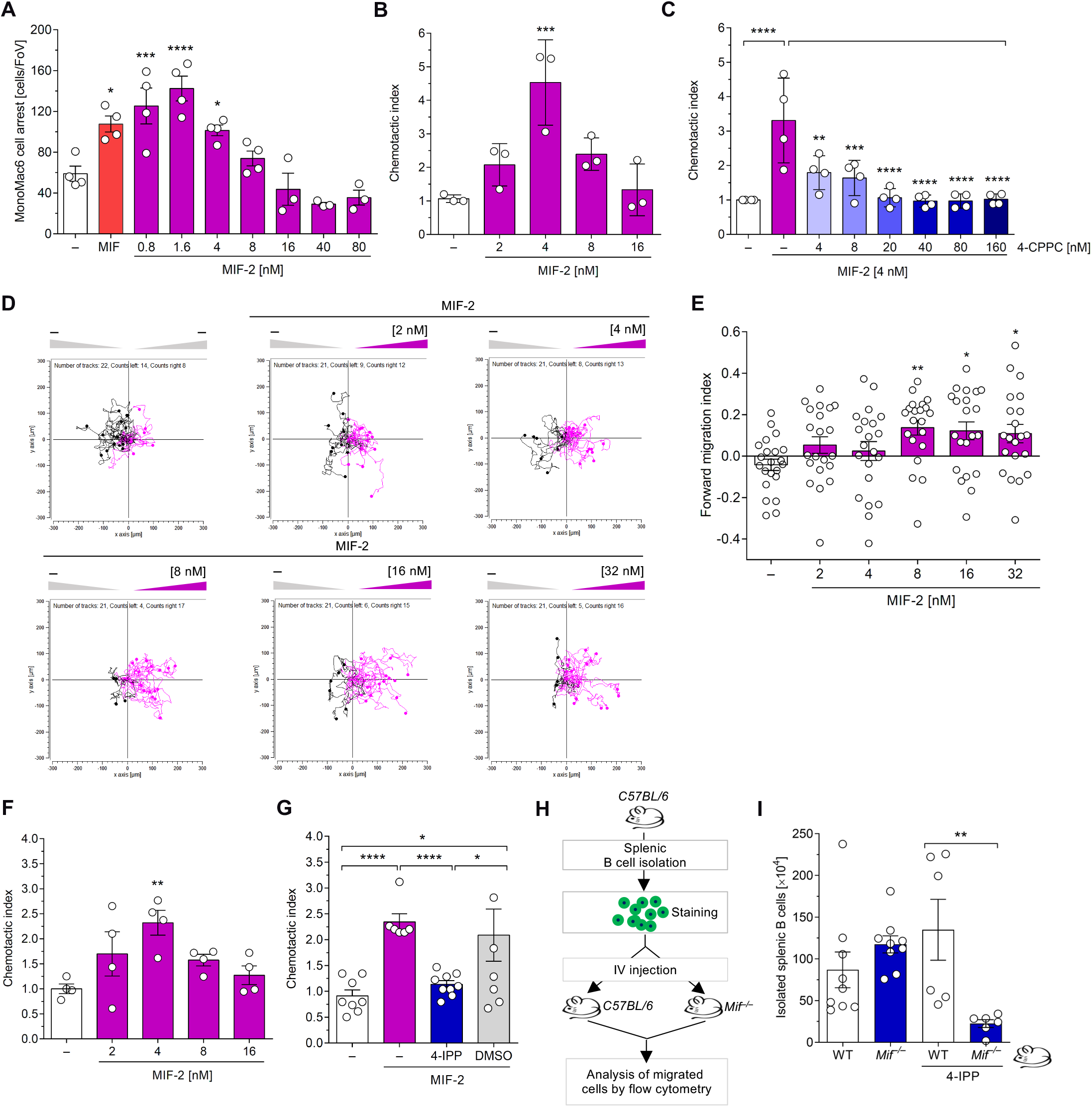
MIF-2 promotes atherogenic leukocyte arrest and chemotaxis. (A) MIF-2 promotes MonoMac6 adhesion on human aortic endothelial cell monolayers (HAoEC) under flow conditions. MIF-2 was tested at the indicated concentrations between 0.8 and 80 nM, with a dose peak observed at 1.6 nM, and its effect compared to that of 16 nM MIF (n = 3-4 biological replicates each). (B, C) MIF-2 triggers monocyte chemotaxis in a 2D Transwell migration set-up and inhibition by 4-CPPC. (B) Human peripheral blood monocytes were subjected to a chemokine gradient of different concentrations of MIF-2 as indicated and relative chemotaxis in relation to random movement in the absence of chemoattractant depicted as chemotactic index. (C) The migratory effect of 4 nM MIF-2 (as tested in (B)) is inhibited by increasing concentrations of the MIF-2-selective inhibitor 4-CPPC (n = 4 biological replicates each). (D, E) MIF-2 triggers monocyte chemotaxis in a 3D migration set-up as assessed by single cell tracking. Dose response for MIF-2 between 2 and 32 nM. (D) Representative experiment demonstrating that MIF-2 elicits (magenta tracks) 3D chemotaxis of human monocytes as assessed by live-microscopic imaging of single-cell migration tracks in x/y direction in µm. The unstimulated control (gray tracks) indicates random motility. The migration tracks of 28–30 randomly selected cells per treatment group were recorded (D) and the forward migration index plotted (E). The experiment shown is one of three independent experiments with monocytes from different donors. (F) MIF-2-induced chemotactic effects on primary B cells via 2D Transwell migration (n = 4 biological replicates). (G) A small selective inhibitor of MIF and MIF-2, 4-IPP was used to verify inhibitory effects on chemotactic properties of MIF-2 (n = 4 biological replicates). (H-I) Effect of MIF-2 on B cell homing *in vivo*. (H) Scheme illustrating the procedural details of the primary B lymphocyte homing assay. Fluorescently labelled primary splenic B cells isolated from wildtype (WT) C57BL/6 mice were i.v.-injected into WT or *Mif ^−/−^* recipient mice and ‘homed’ migrated B cells isolated from target organs quantified by flow cytometry. (I) Quantification of fluorescently labelled primary B lymphocytes homed into spleen according to (H). All values are means ± SD; *p<0.05; **p<0.01; ***p<0.001; ****p<0.0001.

We next asked whether the chemotactic capacity of MIF-2 would extend to other leukocyte populations such as B lymphocytes, which have been implicated in atherogenesis and are known to migrate in response to MIF (16, 52). Notably, MIF-2 elicited the migration of primary splenic B lymphocytes in a concentration-dependent manner, with a maximal chemotactic index of approximately 2.5-fold obtained at a peak concentration of 4 nM, in line with the optimal dose observed for monocyte recruitment responses (Figure 2F). The chemotactic activity of MIF-2 for B cells was confirmed, applying the small molecule compound 4-IPP, an inhibitor of both MIF and MIF-2 (53, 54), which ablated the chemotactic effect of MIF-2, while DMSO-containing solvent control had no effect (Figure 2G). To evaluate the *in vivo* relevance of MIF-2-triggered effects on B lymphocyte trafficking, we studied the migration and homing response of fluorescently labeled splenic B lymphocytes in C57BL/6 mice *in vivo*. To account for effects of MIF versus MIF-2 in a complex *in vivo* setting, wildtype (WT) *versus* global *Mif* gene-deficient (*Mif ^−/−^*) mice were examined (Figure 2H). Additionally, 4-IPP was administered to both mouse groups to pharmacologically block MIF and/or MIF-2. Primary splenic B cells freshly isolated from WT mice were stained with carboxyfluorescein succinimidyl ester (CFSE) and adoptively transferred into WT or *Mif ^−/−^* recipient mice that had been injected with 4-IPP (2.5 mg/kg) or vehicle control 4 h before. Two hours post-injection, CFSE-positive (CFSE^+^) cells were quantified by flow cytometry using cell extracts from spleen, bone marrow (BM), and lymph node (LN) (Figure 2H). Untreated WT and *Mif ^−/−^* recipient mice showed a comparable number of CFSE^+^ B cells homed into the spleen (Figure 2I). This suggested that MIF-2 and/or any other established B-cell chemokine mediated the homing effect. Of note, treatment of mice with the MIF/MIF-2-specific inhibitor 4-IPP led to a marked reduction of B cell homing into the spleen in *Mif*-deleted mice, strongly arguing for a role of MIF-2 in the observed trafficking response of B cells *in vivo* (Figure 2I). A similar, albeit not statistically significant, trend was in BM, while no changes were observed in circulating CFSE^+^ B cell counts or in lymph nodes (LN) (Figure S4A-C). Collectively, these results suggest that MIF-2 promotes the adhesion and migration of leukocytes, and thus qualifies as a novel atypical chemokine.

### MIF-2 interacts with CXCR4 and the MIF-2/CXCR4 axis controls atherogenic activities of MIF-2

Given our findings that MIF-2 promoted the migration of leukocytes in a similar, but seemingly more potent manner, as MIF, we wished to directly compare the migratory capacity of MIF and MIF-2. Primary B lymphocytes were exposed to an optimal chemotactic gradient of MIF-2 (4 nM) in the 3D chemotaxis assay, and their migratory response compared to a set-up in which an optimal concentration of MIF (8 nM) was added to opposite chamber instead of buffer control, so that B cells were placed in the center of competing gradients of MIF-2 and MIF. Of note, B cells showed a pronounced migratory effect towards MIF-2, irrespective of whether MIF or control buffer were placed in the opposite chamber (Figure 3A), confirming that MIF-2 serves as the more potent B cell chemokine than MIF. The sequence alignment of MIF and MIF-2 in Figure 3B illustrates the homology between both MIF proteins across the entire sequence, including for example the presence of the evolutionarily conserved Pro-2 residue, but also indicates significant differences in the putative receptor binding motifs. Most strikingly, the CXCR2 binding signature pseudo-(E)LR motif of MIF is not present in MIF-2, in line with recent findings that MIF but not MIF-2 recruits inflammatory macrophages via CXCR2 in an experimental polymicrobial sepsis model (37). In contrast, most of the residues required for MIF binding to CXCR4 are conserved in MIF-2 (Figure 3B), letting us hypothesize that MIF-2 may also be a ligand of CXCR4.

**Figure 3.**
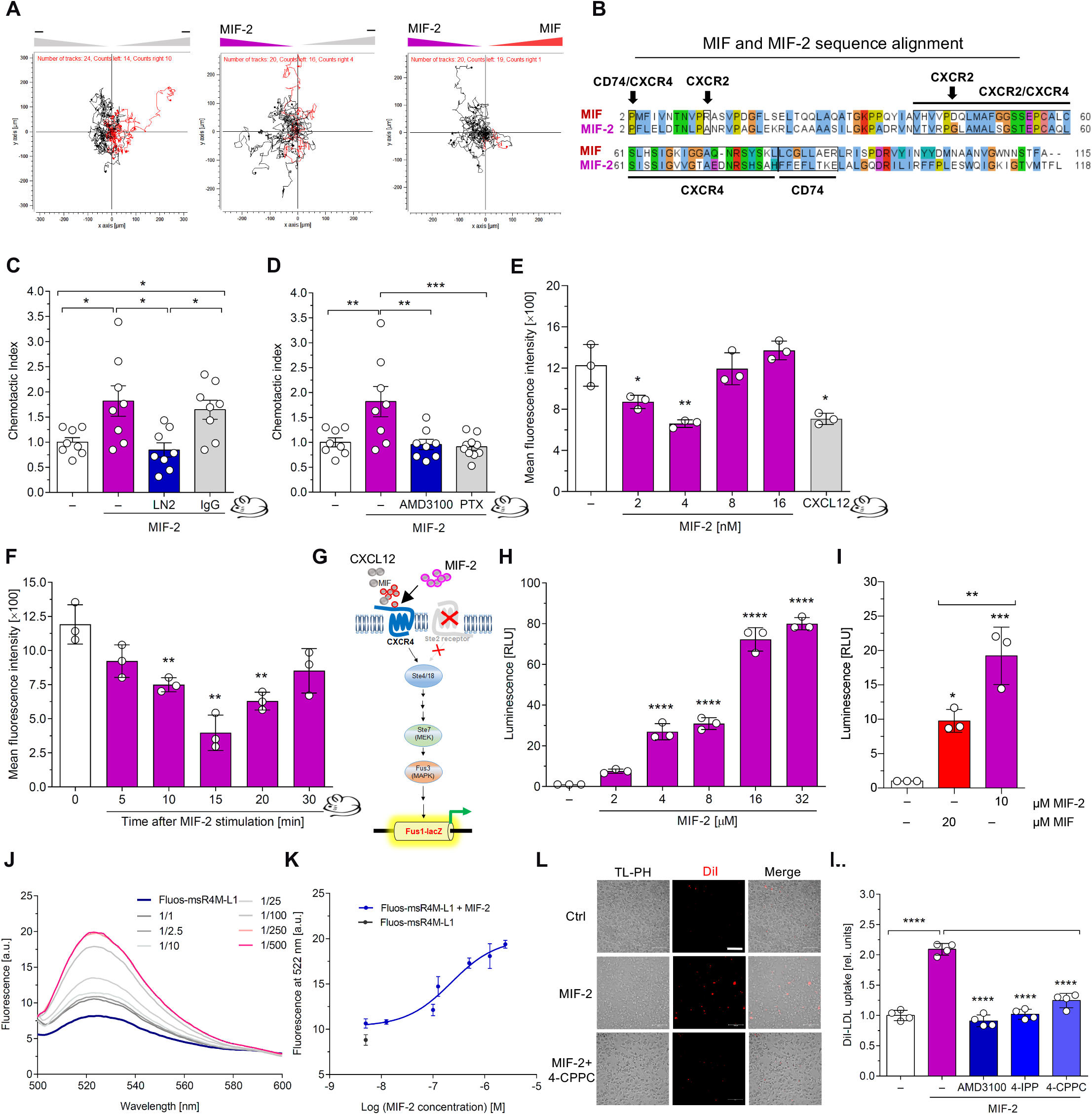
MIF-2 is a more potent chemokine than MIF, is a novel CXCR4 ligand, and promotes foam cell formation in a CXCR4-dependent manner. (A) Direct comparison of the chemotactic potency of MIF and MIF-2 by competing effects on primary B lymphocytes in a 3D chemotaxis assay. Left, negative control with buffer added to both chambers; middle, chemotactic effect of MIF-2 (left chamber) compared to buffer (right chamber); right, direct comparison of the chemotactic potency of MIF-2 (left chamber) with that of MIF (right chamber). The experiment shown is one of three independent experiments performed with B cells from different mice and preparations. (B) Sequence alignment of the amino acid sequences of human MIF-2 and MIF. Sequences were retrieved from the UniProt database and aligned by ClustalW using standard parameters in the Jalview multiple sequence alignment editor desktop application. Residues implicated in the binding of MIF and/or MIF-2 to the different MIF receptors CXCR4, CXCR2, and CD74 are indicated. Identical or homologous residues between MIF and MIF-2 are highlighted by same colors. (C and D) MIF-2 triggers B-cell migration in a CD74- and CXCR4-dependent manner. Transwell migration assay with primary splenic B cells from wildtype (WT) mice. Anti-CD74 antibody (LN_2_) (C) and the CXCR4 antagonist (AMD3100) (D) were applied to test the role of the receptors for MIF-2-mediated chemotaxis of B cells. The relative chemotaxis in relation to random movement in the absence of chemoattractant (MIF-2) is depicted as chemotactic index; the data points shown represent n = 8 biological replicates. (E and F) MIF-2 elicits the internalization of CXCR4 in primary mouse B cells. The internalization of CXCR4 induced by MIF-2 using primary B cells from WT mice was followed in a dose-(E) and time-(F) dependent manner. (G) Scheme summarizing chemokine (MIF, MIF-2, or CXCL12)-induced signaling in the yeast-CXCR4 transformant reporter assay. (H) MIF-2 binds to and signals through human CXCR4 in an *S. cerevisiae* reporter system in a concentration-dependent manner. CXCR4 binding/signaling is read out by LacZ reporter-driven luminescence (n = 3 biological replicates). (I) Same as (H), except that the effect of 10 µM MIF-2 was directly compared with that of 20 µM MIF (n = 3 biological replicates). (J and K) Nanomolar affinity binding of MIF-2 to the CXCR4 surrogate peptide msR4M-L1 as determined by fluorescence spectroscopic titration. Emission spectra of Fluos-msR4M-L1 alone (blue; 5 nM) and with increasing concentrations of MIF-2 at indicated ratios are shown (J; representative titration); binding curve derived from the fluorescence emission at 522 nm (K). The determined app. K_D_ for MIF-2 binding to msR4M-L1 is 180.82 ± 63.07 nM (mean ± SD). (L and M) MIF-2-mediated DiI-LDL uptake in primary human monocyte-derived macrophages is mediated by CXCR4. MIF-2 was applied at a concentration of 80 nM, and the inhibitory effects induced by AMD3100, 4-IPP, and 4-CPPC were evaluated. Representative images of DiI-LDL-positive cells (L) and quantification (three-times-two independent experiments; 9 fields-of-view each) (M). Scale bar, 100 µm. All values are means ± SD; *p<0.05; **p<0.01; ***p<0.001; ****p<0.0001.

Mouse B lymphocytes express high surface levels of CXCR4, as well as the MIF and MIF-2 cognate receptor CD74, and have previously been established as a valuable cell system to study MIF-mediated chemotaxis and receptor internalization responses (16, 24). To begin to study the role of CXCR4 as a potential MIF-2 receptor, we therefore tested the effect of CXCR4 blockade on MIF-2-mediated B-cell chemotaxis and also studied the consequence of CD74 blockade. As expected, and confirming previous data for MIF, a neutralizing antibody against CD74 (anti-CD74, LN-2), but not an isotype control immunoglobulin, blocked MIF-2-elicited B-cell chemotaxis, also serving to further validate this experimental system (Figure 3C). Importantly, co-incubation of B cells with the CXCR4-specific small molecule inhibitor AMD3100/plerixafor, as well as the Giα inhibitor pertussis toxin (PTX), abrogated the MIF-2-mediated B-cell migration response, indicating the involvement of CXCR4 in this process (Figure 3D). As previously found for MIF, the CD74-blocking effect also indicated a co-requirement for CD74.

To obtain additional proof for a possible role of CXCR4 as MIF-2 receptor, we performed flow cytometry-based internalization experiments measuring CXCR4 cell surface levels in primary mouse B cells and determined whether MIF-2 is able to affect CXCR4 internalization. As shown in Figure 3E, MIF-2-treatment dose-dependently triggered the internalization of CXCR4 with a maximal internalization effect seen at 4 nM MIF-2, a potency comparable to that of the CXCR4 cognate ligand CXCL12. Moreover, kinetic studies revealed significant and rapid MIF-2-mediated CXCR4 internalization within 15 min (Figure 3F). Confirming specificity, the effect was observed to be transient as known for chemokine receptor internalization and was again more potent than the effect of MIF, for which maximal internalization of CXCR4 was achieved with at an optimal concentration of 32 nM within 20 min (Figure S5A,B).

To more directly probe the MIF-2/CXCR4 interaction, we next took advantage of a yeast receptor binding assay, in which human CXCR4 is cloned into yeast transformants and coupled to G protein-mediated GPCR-analogous signaling cascade and reporter read-out, previously established to monitor specific CXCR4 binding of CXCL12 and MIF (55) (Figure 3G). Notably, MIF-2 not only triggered CXCR4-mediated signaling in this cell system specific for the transformed human receptor in a concentration-dependent manner (Figure 3H), but also was the more potent CXCR4 agonist, when compared to MIF (Figure 3I). Lastly, we employed a biochemical in vitro assay to directly measure binding of MIF-2 to CXCR4. Relying on a peptide-based soluble CXCR4 mimic msR4M-L1 that we recently identified and which binds MIF with high affinity (56), we performed fluorescence spectroscopic titrations to determine the binding affinity between MIF-2 to msR4M-L1. Remarkably, titration of fluorescently labeled msR4M-L1 (fluos-msR4M-L1) with MIF-2 revealed a strong dose-dependent increase of fluorescence intensity at 522 nm. The derived app. K_D_ was 180 nM (app. K_D_ (fluos-msR4M-L1/MIF-2) = 180.82 ± 63.07 nM), consistent with a high-affinity interaction between MIF-2 and msR4M-L1 (Figure 3J,K). Together with the B-cell chemotaxis, CXCR4 internalization, and yeast-CXCR4 signaling results, these data indicated that MIF-2 serves as a novel and potent non-cognate ligand for CXCR4.

Since MIF, but not CXCL12, was recently shown to specifically promote LDL receptor (LDLR)-mediated foam cell formation in monocyte-derived macrophages in a strictly CXCR4-dependent manner (56, 57), we surmised that this assay could be a valuable cell system to study the role of the MIF-2/CXCR4 axis in atherogenesis. Primary human monocyte-derived macrophages were stimulated with MIF-2 in the presence or absence of 4-IPP, 4-CPPC, or AMD3100, and subsequently exposed to fluorescently labeled native LDL (DiI-LDL) to detect its uptake and thus LDLR-mediated foam cell formation. MIF-2 stimulation led to a marked enhancement of LDL uptake, and this effect was fully inhibited by pretreatment of the cells with all inhibitors (Figures 3L,M), confirming MIF-2-specificity of the foam cell-formation effect and that the MIF-2 effect was CXCR4-dependent (Figure 3L,M). Thus, MIF-2 is a novel ligand of CXCR4 and this axis mediates atherogenesis-relevant activities such as leukocyte migration and foam cell formation.

### *Mif-2* deficiency attenuates the progression of advanced atherosclerosis, lowers body weight, and plasma lipid levels *in vivo*

Since MIF-2 promotes foam cell formation, a critical process observed in advanced stages of atherosclerosis (58, 59), we next studied the impact of *Mif-2-*deficiency on of advanced athero-sclerosis in *Apoe^−/−^* mice. Eight-week-old female *Mif-2^−/−^Apoe^−/−^ versus Apoe^−/−^* littermates were challenged with a HFD for 12 weeks and atherosclerotic plaques were analyzed (Figure 4A). As shown in Figure 4B, *Apoe^−/−^* mice developed pronounced lesions in aortic root with an approximately 3-fold increase of lesion areas compared to those measured in the early-stage feeding model (Figure 1B). Confirming the protective effect in the model of early atherogenesis, global *Mif-2^−/−^* deletion in *Apoe^−/−^* mice attenuated lesion formation, resulting in a significant reduction of lesion size by ∼40% in aortic root (Figure 4B,C), and by ∼80% in aortic arch (Figure 4F), as compared to the control *Apoe^−/−^* mice. *Mif-2*-deficiency also mitigated vascular inflammation as indicated by a reduced lesional macrophage content in *Mif-2^−/−^Apoe^−/−^* compared to control mice (Figure 4D). As expected for an advanced model of atherosclerosis, a marked necrotic core formed as visualized by Masson staining, and notably, this was significantly reduced by *Mif-2* deficiency (Figure 4E), whereas the plaque collagen content remained unaffected (Figure S6A). Surprisingly, the female *Mif-2^−/−^Apoe^−/−^* mice showed a marked reduction in body size at the end of the 12-week-feeding period (Figure 4G) as well as significantly reduced body weight (Figure 4H; Table S1), an observation that was also made after 4.5-week HFD, while the body weights of the mice before start of the diet did not differ between the *Mif-2^−/−^* genotype and the *Mif*-proficient controls (Figure 4H). Similar results were obtained for male cohorts, for which, interestingly, the *Mif-2^−/−^Apoe^−/−^* mice also had slightly but significantly reduced body weights before the start of the HFD (Figure S7A), indicating that in male mice this phenotype is not HFD-dependent, but related to *Mif-2* gene depletion *per se*.

**Figure 4.**
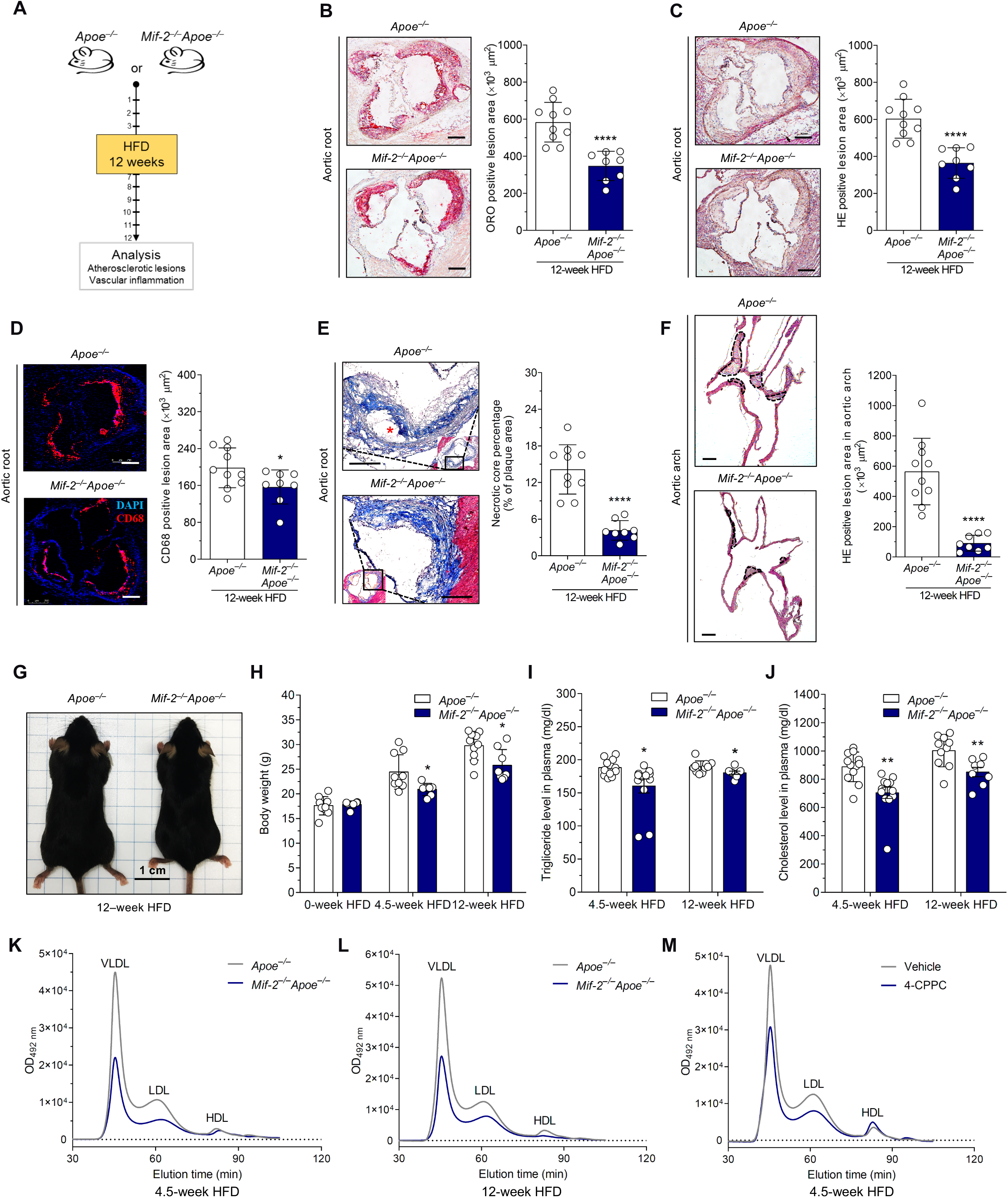
*Mif-2* deficiency ameliorates advanced atherosclerotic lesion progression in hyperlipidemic *Apoe^−/−^* mice. (A) Experimental outline. Female *Apoe^−/−^* and *Mif-2^−/−^Apoe^−/−^* mice were fed with Western-type cholesterol-rich high-fat diet (HFD) for 12 weeks to develop advanced plaques. (B-C) *Mif-2* deficiency reduces the formation of advanced atherosclerotic lesions in aortic root. (B) Representative ORO-stained images of aortic roots in serial frozen sections (8 µm) from *Apoe^−/−^* and *Mif-2^−/−^Apoe^−/−^* mice after 12 weeks of HFD (scale bar: 250 µm) and corresponding quantification (n = 8-10 mice per group; 12 sections per mouse). (C) Same as (B) except that sections were stained with HE. (D) *Mif-2* deficiency reduces lesional macrophage content in aortic root. Representative images (scale bar: 250 µm) and quantification of macrophage area of anti-CD68-stained (red) sections (8 µm) from *Apoe^−/−^* and *Mif-2^−/−^Apoe^−/−^* mice (DAPI, blue). n = 8-10 mice per group; 12 sections per mouse. (E) *Mif-2* deficiency reduces the necrotic core. Necrotic core and collagen content was visualized by Masson staining. A typical classical necrotic core is marked by a red asterisk. n = 8-10 mice per group; scale bar: 200 µm. (F) *Mif-2* deficiency reduces the formation of advanced atherosclerotic lesions in aortic arch. Representative HE-stained images of aortic arch in paraffin sections (4 µm) (scale bar: 200 µm) and quantification results (n = 8-10 mice per group; 8 sections per mouse). (G-H) *Mif-2^−/−^Apoe^−/−^* mice are smaller and lighter than *Apoe^−/−^* mice. Representative gross morphology of a *Apoe^−/−^* mouse compared to a *Mif-2^−/−^ Apoe^−/−^* mouse after 12 weeks of HFD (G) and corresponding body weight results n = 6-10 mice per group (H). (I-J) *Mif-2^−/−^Apoe^−/−^* mice have lower circulating lipid levels than *Apoe^−/−^* mice. Levels of plasma triglycerides (L) and plasma total cholesterol (M) from *Apoe^−/−^* and *Mif-2 ^−/−^Apoe^−/−^* mice fed a HFD for 12 weeks. n = 8-10 mice per group. (K-M) *Mif-2* deficiency or pharmacological blockade reduces atherogenic lipoproteins. Representative HPLC chromatograms of lipo-protein fractions in plasma from indicated mouse cohorts. Female *Apoe^−/−^* and *Mif-2^−/−^Apoe^−/−^* mice after 4.5-week HFD (K); female *Apoe^−/−^* and *Mif-2^−/−^Apoe^−/−^* mice after 12-week HFD (L); male *Apoe^−/−^* mice treated with 4-CPPC *versus* vehicle after 4.5-week HFD (M). All values are means ± SD; *p<0.05; **p<0.01; ***p<0.001; ****p<0.0001.

The measurement of plasma lipids revealed a significant drop of plasma total cholesterol and triglyceride levels by approximately 10%-20% in the *Mif-2^−/−^Apoe^−/−^* mice compared to control after both 4.5- and 12-week-HFD, respectively (Figure 4I,J; Table S1). Moreover, when we analyzed the plasma lipoprotein fractions (very-low-density lipoprotein (VLDL), low-density lipoprotein (LDL), and high-density lipoprotein (HDL)) by HPLC analysis, substantial reductions of VLDL and LDL were observed in *Mif-2-*deficient *Apoe^−/−^* mice compared with *Apoe^−/−^* control mice, as well as in *Apoe^−/−^* mice, in which MIF-2 was pharmacologically inhibited by treatment with 4-CPPC (Figure 4K-M). Taken together, these data suggest that the proatherogenic role of MIF-*2* is not restricted to early stages of atherogenesis but is also important in advanced atherosclerosis and appears to markedly influence plasma lipid and lipoprotein homeostasis.

### MIF-2 promotes hepatosteatosis and regulates SREBP activity to regulate lipogenesis in hepatocytes in a CXCR4- and CD74-dependent manner

Considering that alterations in circulating lipid levels are associated with variations in lipid accumulation in the liver, we next analyzed liver size, liver weight, and hepatic lipid content. The livers of female *Mif-2^−/−^Apoe^−/−^* mice after 12 weeks of HFD were found to be markedly smaller and lighter compared to the livers from *Apoe^−/−^* mice (Figure 5A,B). A similar striking difference was noticed in male mice as well as in control animals that had been on chow diet for 12 weeks, again confirming the significance of *Mif-2* gene depletion *per se* (Figure S7B,C).

**Figure 5.**
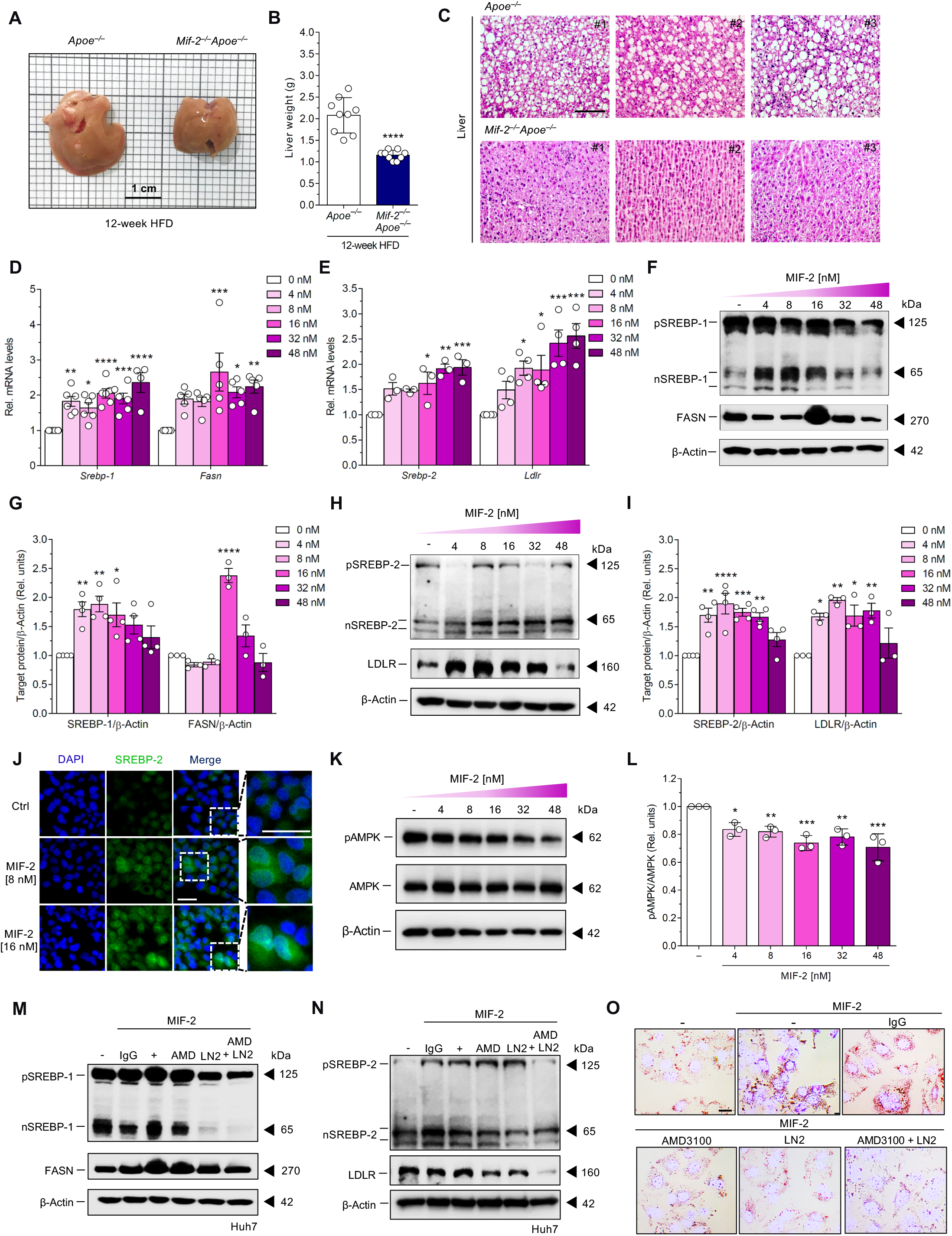
MIF-2 increases hepatosteatosis and elicits SREBP-mediated lipogenesis in hepatocytes. (A-B) Gross morphology of mouse livers (representative images shown) from indicated *Apoe^−/−^* and *Mif-2^−/−^Apoe^−/−^* mice after 12 weeks of Western-type cholesterol-rich high-fat diet (HFD) (A) and corresponding liver weights (B). (C) *Mif-2^−/−^Apoe^−/−^* mice have reduced hepatosteatosis compared to *Apoe^−/−^* mice. Representative HE-stained paraffin liver sections (4 µm) from indicated *Apoe^−/−^* and *Mif-2^−/−^Apoe^−/−^* mice after 12 weeks of HFD (images from n = 3 mice; scale bar: 200 µm). (D and E) Recombinant MIF-2 increases the gene expression of *SREBP* and its lipogenic downstream target transcripts in Huh-7 cells. *Srebp-1, Fasn* (D), *Srebp-2* and *Ldlr* (E) mRNA levels were determined by RT-qPCR following stimulation of Huh-7 cells with increasing concentrations of MIF-2 for 24 h (n = 3-6 biological replicates). (F-I) Same as (D-E) except that protein levels of SREBP precursor (pSREBP) and processed nuclear SREBP (nSREBP) were determined by Western blot. (F-G) MIF-2 upregulates SREBP-1 and FASN protein levels. Representative Western blot (F) and quantification of nSREBP-1 and its target gene FASN (G) (n = 3-4 biological replicates). (H-I) MIF-2 induces SREBP-2 and LDLR protein levels. Representative Western blot (H) and quantification of nSREBP-2 and its target gene LDLR (I) (n = 3-4 biological replicates). β-actin was run as a loading standard. (J) MIF-2 promotes the nuclear translocation of nSREBP2. Stimulation of Huh-7 cells with two different concentrations of MIF-2 compared to buffer control is shown. Immunofluorescent image of SREBP-2 nuclear translocation in in Huh-7 cells stimulated with MIF-2 for 24 h (SREBP2: green; DAPI: blue; right panel shows magnified images; scale bar: 40 µm). (K-L) MIF-2 inhibits the activation of AMPK in Huh-7 hepatocytes. Treatment of Huh-7 cells with rMIF-2 similar to (F) and (H), except that stimulation was for 30 min only and blots were developed for phospho-AMPK and AMPK (K, representative blot; L, quantification of n = 3 independent experiments). β-actin was run as a loading standard. All values are means ± SD; *p<0.05; **p<0.01; ***p<0.001; ****p<0.0001. (M-N) SREBP activation by MIF-2 in Huh-7 cells is dependent on the receptors CD74 and CXCR4. Huh-7 cells were stimulated with 16 nM MIF-2 for 30 min in the presence or absence of the CXCR4 inhibitor AMD3100 (AMD), the CD74-blocking antibody LN2, or combination of both, or immunoglobulin isotype control (IgG) as indicated. (M) Representative Western blot developed for pSREBP-1, nSREBP-1, and the target protein FASN. (N) Representative Western blot developed for pSREBP-2, nSREBP-2, and the target protein LDLR. β-actin was run as a loading control. (O) MIF-2 stimulates LDL uptake in Huh-7 hepatocytes and blockade by CXCR4 and CD74 inhibitors. Native LDL uptake in Huh-7 cells stimulated with 16 nM MIF-2) for 24 h in presence or absence of AMD3100, LN2, or both. An isotype immunoglobulin was used as control (IgG). Cells were stained with ORO to visualize lipid content (scale bar: 25 µm). All values are means ± SD; *p<0.05; **p<0.01; ***p<0.001; ****p<0.0001.

We next determined the hepatic lipid content applying both HE and ORO staining of liver sections. A marked reduction of neutral lipid deposition was detected in *Mif-2^−/−^Apoe^−/−^* mice compared to *Apoe^−/−^* mice (Figure 5C, S8A,B), suggesting a pronounced and causal role for MIF-2 in the liver that may be linked to the regulation of hepatic lipogenesis. To address the possibility that MIF-2 may directly regulate lipid levels in the liver, we determined the effect of MIF-2 on the expression of lipogenic genes in hepatocytes, applying the human hepatocyte cell line Huh-7 and RT-qPCR. We assessed the gene expression of *Srebp-1*, *Srebp-2*, proprotein convertase subtilisin/kexin type 9 (*Pcsk9)*, glucose transporters *Glut1* and *Glut4*, hydroxy-methylglutaryl-coenzyme A reductase (*Hmgcr)*, *Srebp-2*, as well as the scavenger receptor *Cd36* following MIF-2 stimulation of Huh-7 cells for 24 h. MIF-2 treatment resulted in increased mRNA levels of both *Srebp-1* and *Srebp-2* in a dose-dependent manner (Figure 5D,E). In contrast, gene expression of *Glut1*, *Glut4*, *Cd36* (Figure S9A-C) as well as *Hmgcr* and *Psck9* (data not shown) did not change in response to MIF-2 exposure of Huh-7. Furthermore, the gene expression of the MIF (and MIF-2) receptors *Cxcr4* and *Cd74* was not affected by MIF-2 stimulation (Figures S9D,E).

SREBP-1 and SREBP-2 promote the transcription of sterol-regulated genes involved in lipogenesis including fatty acid synthase *(Fasn)* and *Ldlr*, respectively (60, 61). MIF-2 led to an increase in mRNA levels of both *Fasn* and *Ldlr* (Figure 5D,E). Since the activation of SBERP-1 and SREBP-2 is associated with their proteolytic cleavage, we next checked whether MIF-2 would affect SREBP processing. As shown in Figures 5F and G, incubation of Huh-7 cells with increasing concentrations of MIF-2 resulted in a reduction of the SREBP-1 precursor (pSREBP-1) band (∼125 kDa) and in a simultaneous increase in the processed form of SREBP-1 (nSREBP-1) (∼65 kDa) as well as its target gene FASN as revealed by immunoblotting analysis. Similarly, MIF-2 also promoted SREBP-2 processing as well as its key downstream target gene LDLR in a dose-responsive manner in Huh-7 cells (Figures 5H,I). To further verify these findings, we analyzed changes the subcellular localization of the processed form of SREBP-2 following MIF-2 stimulation in Huh-7 cells. In line with the notion that MIF-2 can elicit SREBP-2 processing, cells exposed to MIF-2 showed an enhanced nSREBP-2 accumulation in the nucleus, whereas untreated cells exhibited cytoplasmic localization of the inactive SREBP-2 precursor as shown by immunofluorescence staining (Figure 5J).

Previous studies have shown that induction of the AMPK pathway negatively regulates the processing and transcriptional activity of SREBPs to mitigate hepatosteatosis and atherogenesis (62, 63). We previously found that MIF activates hepatic AMPK, an effect that contributes to its protective effect in fatty liver disease (25). To examine how MIF-2 influences the AMPK pathway in hepatocytes, Huh-7 cells were stimulated with different concentrations of MIF-2 and phosphorylated AMPK (pAMPK) levels, as a readout for AMPK activation, analyzed by immunoblotting. Notably, MIF-2 inhibited pAMPK levels in a concentration-dependent manner with ∼20-30% reduction observed at 16 nM MIF-2 compared to control buffer-treated cells (Figures 5K, L). Thus, MIF-2 appeared to have an opposite effect on the hepatocyte AMPK pathway, in accord with the notion that it promotes SREBP activity, which in turn is down-regulated by AMPK.

### MIF-2-mediated SREBP activation and lipogenic gene expression in hepatocytes is dependent on CXCR4 and CD74

We next aimed to determine whether the observed effects of MIF-2 on hepatocyte lipogenesis are dependent on CD74 and/or CXCR4. Both of these MIF-2 receptors are expressed on Huh-7 cells as revealed by immunofluorescence staining (Figure S10A) and flow cytometry (Figure S10B,C). To investigate an involvement of CXCR4 or CD74 in the observed effects of MIF-2, we performed the MIF-2-elicited SREBP proteolytic activation assay in the presence *versus* absence of inhibitors of CD74 and CXCR4. Blockade of CD74 with the neutralizing LN-2 antibody resulted in a pronounced reduction of processed nSREBP-1 and nSREBP-2 levels, while inhibition of CXCR4 with AMD3100 showed a partial decrease of MIF-2-induced SREBP-1/2 processing and activation (Figures 5M,N). Remarkably, dual inhibition of CXCR4 and CD74 by coincubation with both inhibitors fully abrogated the activating effect of MIF-2 on the cleavage of SREBP-1 and SREBP-2 as well as the expression of the downstream target proteins FASN and LDLR (Figures 5M,N), providing plausible evidence that the MIF-2-mediated lipogenic effects in Huh-7 cells are dependent on CXCR4/CD74 axis.

To further examine potential functional consequences of this pathway on hepato-steatosis and lipoprotein turnover, we measured the effect of MIF-2 on hepatic LDL uptake and the contribution of CXCR4 and CD74 to this process. Huh-7 cells were pretreated with AMD3100, LN-2 antibody, or control IgG, alone or in combination, stimulated with MIF-2 and exposed to LDL. ORO staining revealed that MIF-2 stimulation led to markedly enhanced LDL uptake, an effect that was substantially attenuated by CXCR4 or CD74 inhibition (Figure 5O). Noteworthy, coincubation with AMD3100 and LN resulted in a maximal inhibitory effect, together suggesting that MIF-2-driven LDL uptake may contribute to hepatic lipid accumulation and that this effect is mediated by both CD74 and CXCR4.

### MIF-2 expression is strongly upregulated in unstable human atherosclerotic plaques

Our *in vitro* data and experimental *in vivo* studies applying Mif-2-deficient *Apoe^−/−^* mice suggest that MIF-2 is a potent chemokine that promotes vascular inflammation and atherogenesis both in the early and advanced stage. Moreover, MIF-2 enhances foam cell formation, and surprisingly increases hepatosteatosis and lipogenic gene expression in hepatocytes. Atherosclerotic lesions of *Mif-2^−/−^Apoe^−/−^* mice also displayed markedly reduced necrotic core regions, altogether suggesting that MIF-2 may have a particularly critical role in advanced atherosclerosis.

To further test this notion and explore its translational significance, we analyzed the expression and potential deregulation of MIF-2 in human stable and unstable carotid artery plaques collected from patients undergoing endarterectomy. The plaque phenotype (stable/unstable) was assigned based on the morphology following the American Heart Association (AHA) classification (64) as well as fibrous cap thickness (below or above 200 µm) according to Redgrave et al. (65).

Scouting data obtained by DAB (3,3’-diaminobenzidine)-based immunohistochemical (IHC) staining of MIF-2 protein with an established anti-MIF-2 antibody in paraffin-embedded plaque sections suggested that MIF-2 protein is abundantly present in unstable plaque tissue, whereas markedly less MIF-2 immunopositivity was detected in stable plaque tissue (Figure 6A). For control, sections from healthy vessel tissue were stained and MIF-2 positivity was found to be light and confined to a thin area around the endothelial lining, and control stainings without primary antibody were fully devoid of immunopositivity. This finding encouraged us to more comprehensively analyze and quantify MIF-2 expression in plaque tissue by RT-qPCR. Comparing tissues specimens from stable *versus* unstable plaques (15 patients per group), MIF-2 expression was found to be significantly higher in unstable plaques (Figure 6B). As upregulated MIF expression levels do not differ between stable and unstable CEA plaques (56), this finding is in accord with the notion that MIF-2 may critically contribute to plaque destabilization and rupture.

**Figure 6.**
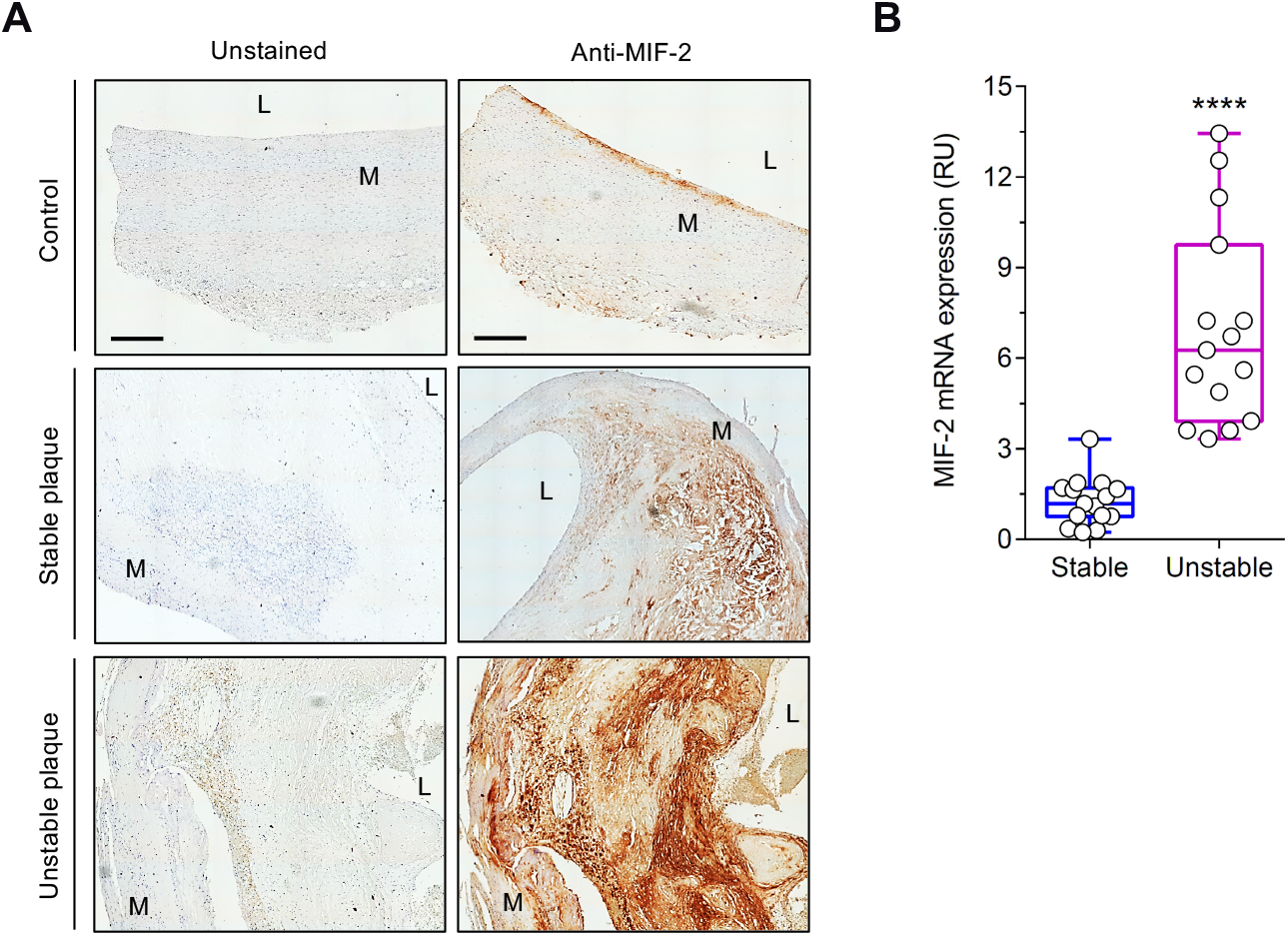
MIF-2 is highly expressed in unstable human carotid plaques. (A) Immunohistochemical staining showing MIF-2 positivity in stable and unstable carotid plaque obtained from carotid endarterectomy patients and comparison to a healthy vessel. Representative DAB (3, 3’-diaminobenzidine) staining images from paraffin-embedded plaque sections developed with an anti-MIF-2 antibody (right panel). Scale bar: 500 µm. The images shown are representative of 5-6 vessels in each group. Left panel, negative control without primary antibody. (B) MIF-2 mRNA expression in stable versus unstable carotid palques as determined by RT-qPCR. n = 15 patients per group. The values are means ± SD; ****p<0.0001.

## DISCUSSION

The present study is the first to identify macrophage migration inhibitor factor-2/D-dopachrome tautomerase (MIF-2/D-DT) as an atypical chemokine and non-cognate ligand of CXCR4 that promotes atherogenesis and vascular inflammation. In this capacity, MIF-2 does not just share its pro-atherogenic properties with MIF, but surprisingly, and contrary to MIF, our data suggest that it enhances circulating atherogenic lipids, hepatosteatosis, and lipogenic pathways in hepatocytes. In line with this vascular and hepatic/lipid phenotype, MIF-2 was found to be markedly upregulated in unstable human carotid plaques, together suggesting that MIF-2 is a novel atherogenic mediator that may regulate lipogenesis and vascular inflammation in cardiovascular diseases.

Using mouse models of both early and advanced atherosclerosis, we found that global deletion of the *Mif-2* gene in *Apoe^−/−^* mice protected against HFD-induced atherogenesis and led to a marked reduction in lesion formation, vascular inflammation, and circulating inflammatory cytokines. These results suggest that MIF-2 can be classified as a novel pro-atherogenic cytokine. The observed lesional phenotype differs from that seen for the MIF-2 homolog MIF. We previously showed that *Mif^−/−^Apoe^−/−^* mice fed with a HFD for 16 and 24 weeks displayed a regiospecific phenotype, resulting in a significant reduction in plaque size in the brachiocephalic artery and in the abdominal aorta, whereas no changes were observed in aortic root, aortic arch, and thoracic aorta when compared with *Apoe^−/−^* control mice (52). In contrast, when analyzing the atherosclerotic lesions in *Mif-2^−/−^Apoe^−/−^* mice after 4.5 and 12 HFD, we found a marked decrease of lesions in aortic root and arch compared with *Apoe^−/−^* mice, with a small decrease noted for circulating monocytes. Results from studies of *Mif-2^−/−^ Apoe^−/−^* mice on a 24-week HFD treatment regimen are not available yet, but we would predict a similar protective phenotype for the *Mif-2*-deficient mice across the entire vascular bed. The protective effect of global *Mif-2* gene deletion was confirmed in a pharmacological model, applying the MIF-2-selective small molecule inhibitor 4-CPPC (32, 51). Consistently, similar results were obtained in 4-CPPC-injected *Apoe^−/−^* mice after 4.5-week HFD. This finding also rules out any gene compensation effects that may be seen in global gene knockout models. Together, the *in vivo* atherosclerosis experiments indicate MIF and MIF-2 exhibit similar but also distinct vascular functions and mechanisms in driving the progression of atherosclerosis. This overall conclusion regarding the atherogenic effects of MIF-2 also was underpinned the atherogenic *in vitro* studies. Similar to MIF (16, 56), MIF-2 was found to dose-dependently enhance the chemotactic migration and endothelial arrest of monocytes as well as B-cell chemotaxis and B-cell homing *in vivo*. However, the different dose response curves and a direct comparison between the chemotactic potency of MIF and MIF-2 strongly suggested that MIF-2 may be considered the more potent chemokine compared with MIF.

Similar to MIF, MIF-2 has been described as a high-affinity ligand for CD74 (30). Since cell surface CD74 has a short cytosolic segment, it initiates intracellular signaling either by activation of the signaling co-receptor CD44 (21) or by regulated intramembrane proteolysis, although the later pathway had not examined outside of B lymphocytes (66, 67). Indeed, MIF-2 interacts with CD74 to trigger the activation of the Erk1/2 MAP kinase pathway via CD44 (30). However, Ishimoto *et al.* also reported that the MIF-2/CD74 axis can drive the expression of IL-6 independently of CD44 in preadipocytes (43), suggesting the presence of an alternative pathway such as regulated intramembrane proteolysis. Interestingly, cell biological and biochemical studies have suggested that CD74 can also form heterodimers with the MIF chemokine receptors including CXCR4 (15, 23, 24). Moreover, a recent study by Song *et al.* exploring the role of MIF-2 in lung epithelial cells suggested that MIF-2-induced PI3K/AKT signaling is mediated by CXCR7 but not by CD74 or CXCR4 (33, 34), while Tilstam *et al.* demonstrated that MIF but not MIF-2 recruits inflammatory macrophages in a polymicrobial sepsis model via CXCR2 (37). Applying an array of biochemical and immunological methods, in our current study, we identify CXCR4 as a novel receptor for MIF-2 and provide evidence that the MIF-2/CXCR4 axis is critical in conveying the atherogenic activities of MIF-2.

When comparing the amino acid sequences of MIF and MIF-2, we noted the presence of conserved motifs that may be important for the interaction between MIF and CXCR4. In contrast, MIF-2 lacks the so-called pseudo(E)LR motif shown to contribute to the MIF/CXCR2 interaction and as discussed above, MIF-2 is unable to recruit inflammatory macrophages via CXCR2 (37). Combining data from internalization, yeast-CXCR4 transformant, and titration experiments, we demonstrated that MIF-2 interacts with CXCR4 with nanomolar affinity and in a dose-dependent manner to elicit cell responses in model cells and leukocytes. These findings were supported by inhibitor studies showing that pharmacological blockade of CXCR4 led to a reduction or abrogation of MIF-2-mediated effects, suggesting that MIF-2 serves as a novel non-cognate ligand of CXCR4. Importantly, the effects mediated by MIF-2 via CXCR4 appear to be more pronounced than those elicited by MIF via the same receptor in both the internalization and yeast-CXCR4 transformant assays, suggesting that MIF-2 may be the more potent ligand for CXCR4 with respect to functional signal transduction than MIF. This notion would be in line with our observation that MIF-2 behaved as the more potent chemokine, at least in the cell systems tested in our study. The obtained apparent affinity constant for the MIF-2/msR4M-L1 interaction in the performed fluorescence titration experiment confirmed high-affinity binding between MIF-2 and CXCR4, but also was approximately 6-fold higher than that previously determined for MIF (180 versus 30 nM) (56). However, this apparent difference is within the range of fluctuations seen between biophysical measurements and for example could be due to different buffer conditions used. Maybe more importantly, msR4M-L1 was designed to specifically interact with MIF and thus is not ‘optimized’ for MIF-2 binding.

MIF-2 is expressed in various tissues and particularly high levels have been noted in the liver (30, 68). However, the role of MIF-2 in this organ has not been explored. A novel and particularly intriguing finding of our study is that *Mif-2^−/−^Apoe^−/−^* mice exhibit a liver phenotype after HFD compared to Mif-2-proficient *Apoe^−/−^* mice that is associated with a significant decrease in total plasma cholesterol and triglyceride levels and markedly reduced hepatosteatosis, suggesting the involvement of MIF-2 in lipid metabolism. Since the biosynthesis of cholesterols, fatty acids and triglycerides are regulated by SREBPs (49), we studied the role of MIF-2 in SREBP activation in hepatocytes. Of note, previous studies reported that genetic depletion of *Srebp-1 in vivo* was associated with a reduction in hepatic fatty acid production, whereas overexpression of Srebp-1 or Srebp-2 in mice promoted steatosis with an increase in plasma triglycerides and cholesterol levels (69, 70). In contrast to SREBP-1a, SREBP-1c is the most predominant isoform expressed in the liver (70). Therefore, it is conceivable that its regulation and activation is controlled by MIF-2. The activation of SREBPs is initiated by their proteolytic processing that is regulated by various factors including sterol, oxysterol and insulin. Here, we provide evidence that MIF-2 acts as a novel regulator of SREBP activation and processing in hepatocytes. Our results in the human hepatocyte cell line Huh-7 indicate that MIF-2 dose-dependently promotes the activation both SREBP-1 and - 2 and the expression of their target genes by inducing SREBP proteolytic cleavage and nuclear translocation. The cleavage and subsequent translocation of the SREBPs is mediated by the well-conserved SCAP-S1P-S2P axis (50, 71). However, the upstream signaling components involved in the regulation and activation of this axis remain unexplored.

One important observation in our study is that the individual or simultaneous blockade of either CXCR4 or CD74 abolished MIF-2-mediated SREBP proteolytic activation, suggesting that these receptors are upstream of the proteolytic process. We would speculate that CXCR4 and CD74 either operate via receptor complex formation or signaling crosstalk between CXCR4- and CD74-induced pathways. Of note, Kim *et al.* demonstrated that the CXCL12/CXCR4 axis is involved in SREBP-1-mediated FASN expression by enhancing the cleavage and nuclear translocation of SREBP-1 to modulate the upregulation of FASN in cancer cells (72). Unlike the PI3K/AKT pathway, which promotes lipogenesis, AMPK activation negatively regulates the expression of SREBP-1c in mouse livers and rat hepatoma cells (73, 74). Furthermore, polyphenol-induced AMPK signaling suppresses the proteolytic processing of both SREBP-1 and SREBP-2 to attenuate lipid biosynthesis in the liver, resulting in reduced hepatic steatosis (62). In light of these observations, we assessed whether MIF-2 would modulate the AMPK pathway in liver cells. Indeed, MIF-2 stimulation of Huh-7 cells was associated with an attenuation of AMPK phosphorylation. This suggests that MIF-2 could be a regulator of lipid homeostasis that affects SREBP activity through suppressing the AMPK pathway in hepatocytes. This notion would be in line with previous studies that have suggested a link between MIF-2 and AMPK signaling in adipocytes and cardiomyocytes (42, 45).

The role of MIF and MIF-2 in hepatic steatosis and liver lipid metabolism appear to be oppositional. Previous data indicated that MIF has a hepatoprotective effect in steatosis and fibrosis by promoting the activation of CD74/AMPK pathway in hepatocytes and *Mif^−/−^* mice on a non-Western-type HFD showed enhanced lipid accumulation in the liver, which was associated with an upregulation of *Srebp-1* and *Fasn* mRNA levels (25, 75). In contrast, while the specific high-fat diet was different (triglyceride-rich *versus* Western-type), our *in vivo* and *in vitro* results suggest a differential – oppositional – role of MIF and MIF-2 in the liver, particularly regarding the regulation of lipid metabolism.

Since the pro-lipogenic role of MIF-2 could contribute to an exacerbation of the vascular effects of MIF-2 and since our data obtained in the *in vitro* and *in vivo* atherogenesis models pointed to a potentially particular role of MIF-2 in advanced atherogenesis, we lastly analyzed MIF-2 expression levels in human atherosclerotic plaque tissue, comparing stable and unstable plaques from patients undergoing CEA, several of which also suffer from diabetic and metabolic co-morbidities. MIF-2 was found to be elevated in atherosclerotic plaques compared to healthy vessel tissue, and notably, MIF-2 expression was strongly upregulated in unstable plaques compared to stable plaques. This observation is striking, as MIF expression levels, while also generally upregulated in CEA plaques compared to healthy vessels, did not differ between stable and unstable plaque tissue (56). This could be suggestive of differential roles of both MIF proteins in different stages of atherosclerosis progression.

In summary, our study extends the significance of the MIF protein family in the pathogenesis of atherosclerosis and identifies MIF-2 as a novel proatherogenic atypical chemokine. MIF-2 may be the more potent atherogenic chemokine for monocytes and B cells compared to its homolog MIF, and in contrast MIF, also appears to contribute to lipid accumulation in the liver. This work also gives initial insight into the underlying mechanism of how MIF-2 activates SREBPs in hepatocytes to promote lipogenesis. Overall, these data also contribute to our understanding of how vascular inflammation and adverse hepatic phenotypes could be linked in atherosclerotic diseases.

## ACKNOWLEDGMENTS

This work was supported by Deutsche Forschungsgemeinschaft (DFG) grant SFB1123-A3 to J.B. and A.K., DFG INST 409/209-1 FUGG to J.B., SFB1123-A1 to C.W., SFB1123-Z1 to R.T.A.M., and SFB1123-B5 to L.M., and by DFG under Germany’s Excellence Strategy within the framework of the Munich Cluster for Systems Neurology (EXC 2145 SyNergy—ID 390857198) to J.B. and C.W. C.Z. acknowledge support by the China Scholarship Council (CSC) fellowship program CSC/LMU. R.B. is supported by the US National Institutes of Health. We are thankful to Dr. Yaw Asare and Dr. Sijia Wang for valuable discussions and to Simona Gerra and Kathleen Hille for technical assistance with rMIF-2 preparations and peptide synthesis and purification, respectively.

## DECLARATION OF INTERESTS

J.B., R.B., and C.W. are co-inventors of patents covering anti-MIF and anti-cytokine strategies for inflammatory and cardiovascular diseases. C.Ko., A.K., O.E., and J.B. are co-inventors of a patent application covering MIF-binding CXCR4 ectodomain mimics for inflammatory and cardiovascular diseases. The remaining authors declare no competing interests.

## MATERIALS AND METHODS

### Atherosclerotic mouse models and treatment

All mouse experiments were approved by the Animal Care and Use Committee of the local authorities and performed in accordance with the animal protection representative at the Center for Stroke and Dementia Research (CSD). *Apoe^−/−^* mice with C57BL/6 background were initially obtained from Charles River Laboratories and backcrossed within the CSD animal facility before use. Even though there were no methods of statistics used to predetermine sample size, G power analysis was applied in this study to validate enough number of mice in all cohorts. To generate *Mif-2^−/−^Apoe^−/−^* mice, *Mif-2^+/–^* and atherogenesis-prone *Apoe^−/−^* mice were cross-bred and housed at the CSD. Genetic mouse experiments were conducted on eight-week-old female and male *Mif-2^−/−^Apoe^−/−^* mice and *Apoe^−/−^* mice. The distribution of these mice was not random and researchers were not blinded by allocation during experiments and outcome assessment. For pharmacological blockade of MIF-2, 4-CPPC (Aurora Fine Chemicals LLC) was administered to *Apoe^−/−^* mice. Eight-week-old male *Apoe^−/−^* mice were divided into two groups of 11 mice each at random. The experimental group was intraperitoneally administrated with 50 µg 4-CPPC dissolved in physiological saline (0.9% NaCl) every other day for 4.5 weeks; the control group was injected with saline at the same time intervals. The 4-CPPC-injection showed no toxicity and no other side effects on mice. The non-selective MIF family cytokine inhibitor 4-IPP was obtained from Tocris Bioscience.

All mice were housed under a 12 h light/dark cycle with *ad libitum* access to food and water. At the age of 8 weeks, mice were fed with a Western-type high-fat diet (HFD) containing 0.21% cholesterol (ssniff Spezialdiäten GmbH) for 4.5 or 12 weeks. Early-to-intermediate atherosclerotic lesions develop in these mouse models (76). At the prospective endpoint of the planned experiment, mice were assigned to isoflurane and midazolam/medetomidine/ fentanyl (MMF) anesthesia, body weight was measured, and blood was collected by cardiac puncture for routine immune cell counts and lipid analysis. After that, mice were transcardially perfused with saline, hearts and proximal aortas were prepared and fixed for quantitative plaque analysis and vessel morphometry, and other organs such as spleens, livers and adipose tissues were stored in –80°C.

### Cell lines

MonoMac-6 cells (a human monocytic cell line derived from a patient with relapsed acute monocytic leukemia (77)) were cultured in RPMI 1640 GlutaMAx medium (GIBCO) supplemented with 10% fetal calf serum (FCS) (GIBCO), 1x non-essential amino acids (NEAA), and 1% penicillin/streptomycin (P/S) (GIBCO). Human aortic endothelial cells (HAoECs; initially isolated from healthy human aorta) were acquired from PromoCell GmbH. After thawing, cells were cultured in endothelial cell growth medium (ECGM) from the same company. Cells were plated on cell culture plates/dishes coated with collagen (Biochrom AG). NIH/3T3 mouse embryonic fibroblasts and the human hepatocellular carcinoma cell line Huh-7 were purchased from Deutsche Sammlung für Mikrobiologie und Zellkulturen (DSMZ). They were cultured in DMEM-GlutaMAX (GIBCO) complemented with 10% FCS and 1% P/S. All cells were kept in a humidified CO_2_ incubator at 37°C. For sub-cultivation, cells were split 2 to 3 times per week with a 1:3-1:5 split ratio.

### Primary cells

The isolation procedures of peripheral blood mononuclear cells (PBMCs) and monocytes were approved by the local ethics committee of the LMU Munich University and all experiments were carried out in accordance with its guidelines. For purification and differentiation of primary human monocytes, PBMCs were isolated as previously described (78). In brief, peripheral blood was obtained from healthy donors and mixed gently with pre-warmed phosphate buffered saline (PBS) (Invitrogen) in a ratio of 1:1. The layer of PBMCs was separated by density gradient centrifugation using Ficoll-Paque Plus density gradient media (GE Healthcare). After centrifugation, cells in the interphase were carefully aspirated and subsequently washed with PBS. The cell pellet was resuspended in RPMI 1640 medium and PBMCs were maintained at 37°C in a humidified atmosphere of 5% CO_2_. Primary human monocytes were then purified by negative depletion using Pan Monocytes Isolation Kit, human (Miltenyi Biotec) following the manufacturer’s protocol. The purity of isolated monocytes was analyzed by flow cytometry using an anti-CD14 antibody (Miltenyi Biotec). The purified monocytes were ready-to-be-used for the Transwell migration and 3D chemotaxis assays as described below.

The differentiation of isolated monocytes into macrophages was performed in RPMI 1640 GlutaMAx medium with 10% FCS and 1% P/S containing 100 ng/mL macrophage colony-stimulating factor (M-CSF) (PeproTech) for 7 days. At the seventh day, the medium was replaced with fresh medium, and macrophages were subjected to the DiI-LDL uptake assay.

For primary murine splenic B-cell isolation, spleens were placed immediately in Eppendorf tubes containing pre-cold PBS on ice and then transferred to a 40 µm cell strainer in a small dish. Tissue was homogenized using 10 mL syringes gently and collected in a 15 mL falcon tube. The cell pellet was kept after centrifugation and red blood cells (RBCs) were removed using 3 mL RBC lysis buffer (Pierce) at room temperature (RT) for 3 min, and then 27 mL PBS was added to terminate the reaction. After washing, all cells were counted by a TC20™ Automated Cell Counter (Bio-Rad). Splenic B cells were purified by depletion of the magnetically labeled cells using Pan B Cell Isolation Kit, Mouse (Miltenyi Biotec), according to the manufacturer’s protocol, and subsequently suspended in RPMI 1640 medium with 10% FCS and 1% P/S ready to be applied in the Transwell migration and 3D chemotaxis assay.

### Tissue preparation and lesion analysis

The heart tissues saved for plaque analysis were embedded in Tissue-Tek optimum cutting temperature (O.C.T.) compound and directly fresh-frozen on dry ice in preparation for sectioning. After the block was trimmed, serial eight-µm thick frozen sections were arranged for Oil Red O (ORO), hematoxylin and eosin (HE) staining, and subsequent quantification of other plaque components, such as immune cells, collagen and necrotic core. The lipid content in aortic root was stained with 0.5% ORO solution in propylene glycol (Sigma-Aldrich) at 37°C for 45 min and nuclei were lightly stained with hematoxylin at RT for 1 min. The lesion area was alternatively stained with hematoxylin (Sigma-Aldrich) for 10 min and then counterstained with eosin (Sigma-Aldrich) for 30 sec. The macrophage content in plaques of the aortic root was visualized by a rat anti-CD68 antibody (1:100) in combination with a Cy5-conjugated secondary antibody (1:300). Meanwhile nuclei were stained with DAPI. Previously isolated and trimmed aortic arch was fixed in 1% paraformaldehyd (PFA) overnight and transferred into PBS on the day before dehydration. After immersed completely, samples were embedded in paraffin. Molded blocks can be stored at RT or be used for direct sectioning. Four µm paraffin sections including three main branches were cut and HE-stained for plaque measurement. In addition, collagen and necrotic core were stained in consonance with the manufacturer’s procedures of trichrome stain (Masson) kit. Nuclei stains black, cytoplasm and muscle fibers stain red, whereas collagen displays blue coloration. Images were acquired with a Leica DMi8 fluorescence microscope (Leica Microsystems), and signals were quantified using computer-assisted image analysis software (ImageJ).

### Inflammatory cytokine array

Plasma cytokine and chemokine profiles from *Mif-2^−/−^Apoe^−/−^* mice and *Apoe^−/−^* mice were mapped by capitalizing on a membrane-based mouse cytokine array panel A (R&D Systems) following the standard instructions. Samples were constituted by placing mouse plasma in a 1:10 dilution of array buffer, mixed with a detection antibody cocktail (1:100), and incubated at RT for 1 h. Then pre-blocked membranes were covered with reconstituted samples at 4°C overnight. The second day, membranes were rinsed followed by 30 min of incubation with the diluted streptavidin-horseradish peroxidase (HRP) solution (1:2000), and then visualized with chemi-reagent mix and developed by an Odyssey® Fc imager (LiCor) for 2 min, 10 min and 1 h, respectively. All measurements in this assay were conducted in duplicates. The mean pixel density of each pair of duplicate spots was quantified by ImageJ and is represented as the relative level of corresponding cytokine.

### Gene expression analysis by RT-qPCR

#### Analysis in Huh-7 hepatocytes and mouse heart tissue

For all experiments involving gene expression analysis, total RNA from cultured cells or tissues was isolated, and then concentrations of RNA samples were measured by Nanodrop spectrophotometer (ThermoFisher Scientific). The real-time quantitative PCR (RT-qPCR) analysis was carried out by using the 2x SensiMix PLUS SYBR No-ROX Kit (Bioline) in a Rotor-Gene 6000 (Corbett) and employing specific primers, which were purchased from Eurofins Genomics. Raw data were acquired via the Rotor-Gene 6000 Series 1.7 software (Corbett) and relative mRNA levels were calculated by using the ΔΔC_t_ method with *β-actin* as a housekeeping gene. PCR primers are detailed in Table S2.

For analysis in cultured cells, 4 x 10^5^ Huh-7 hepatocytes were seeded in the 12-well plate on the day before starting the experiment and then starved with DMEM containing 2% FCS overnight. On the third day, Huh-7 cells were stimulated with different concentrations of purified recombinant MIF-2 (i.e. 0 nM, 4 nM, 8 nM, 16 nM, 32 nM, 48 nM) for 24 h. Afterwards, cells were lysed and RNA was isolated using TRIzol™ Reagent (ThermoFisher Scientific) following the manufacturer’s protocol. For murine tissues, half of the mouse heart was collected and flash-frozen on dry ice and then transferred to a 40 µm cell strainer (Corning). The tissue was cut into small pieces and ground thoroughly using a pipette tip. Genomic DNA was removed from the lysate and total RNA was purified through washing and elution using the RNA/Protein Purification Plus kit (Norgen Biotek) according to the manufacturer’s protocol.

#### Analysis in human carotid plaque tissue

Homogenization of human carotid artery plaque tissue was performed in 700 µL Qiazol lysis reagent and total RNA was isolated using the miRNeasy Mini Kit (Qiagen) according to the manufactureŕs instruction. RNA concentration and purity were assessed using NanoDrop. RIN number was assessed using the RNA Screen Tape (Agilent, USA) in the Agilent TapeStation 4200, and only samples with a RIN > 5 and DV200 of >60% were included for further analysis. Next, cDNA synthesis was performed using the High-Capacity-RNA-to-cDNA Kit (Applied Biosystems, USA) according to the manufacturer’s instructions. Quantitative real-time PCR was performed in 96 well plates with a QuantStudio3 Cycler (Applied Biosystems), using TaqMan Gene Expression Assays (ThermoFisher) for detection of RPLPO (housekeeping) and MIF-2/D-DT.

### Signaling pathway experiments and Western blot analysis

For all experiments involving protein analysis using Western blot, protein samples were denatured at 95°C for 10 min. For immunoblot analysis, 10-20 μL protein/lane (corresponding to around 10-20 μg total protein) was loaded into an SDS-PAGE gel, using gels of different acrylamide percentage according to the molecular weight of target proteins. Electrophoresis was carried out and proteins electro-transferred to a PVDF membrane. After blocking with 5% BSA-TBST, membranes were incubated with primary antibodies diluted in 5% BSA-TBST, as follows: anti-SREBP-1 (1:500), anti-SREBP-2 (1:1000), anti-FASN (1:500), anti-LDLR (1:500), anti-MIF-2 (1:1000), anti-AMPK (1:1000), anti-pAMPK (1:1000). After overnight incubation with primary antibody, the membranes were rinsed in 1x TBST for three times and incubated with goat anti-mouse or rabbit HRP-linked secondary antibody (1:10,000). Following thorough washing, the immunoblot was developed applying chemiluminescence and eventually visualized by an Odyssey® Fc imager.

To study the SREBP signaling pathway *in vitro*, 1 x 10^5^ Huh-7 cells were seeded in a 24-well plate on the day before starting the experiment and then starved with DMEM including 2% FCS overnight. On the third day, Huh-7 cells were stimulated with different concentrations of MIF-2 (i.e. 0 nM, 4 nM, 8 nM, 16 nM, 32 nM, 48 nM) in 2% FCS DMEM for 24 h. Moreover, Huh-7 cells were also incubated by 8 nM MIF-2 with or without IgG, AMD3100 and LN-2 for 24 h to evaluate the blocking effects of receptors. After stimulation, Huh-7 cells were lysed in 1 x lysis buffer with dithiothreitol (DTT) and boiled for immunoblotting analysis. As for the AMPK signaling pathway analysis, Huh-7 cells were stimulated with different concentrations of MIF-2 in 0.05% FCS DMEM for 30 min and samples processed as above.

### Immunofluorescence analysis of nuclear SREBP

1 x 10^4^ Huh-7 cells were seeded in a 24-well plate on the day before starting the experiment and starved with 2% FCS DMEM overnight. On the third day, Huh-7 cells were stimulated with two concentrations of MIF-2 (8 and 16 nM) in 2% FCS DMEM for 24 h. After stimulation, Huh-7 cells were fixed with 4% PFA and permeabilized with 0.02% Triton X-100 in PBS. Cells were washed with PBS, blocked with 1% BSA and 5% goat serum in PBS and then incubated with mouse anti-SREBP-1 (1:100) and mouse anti-SREBP-2 (1:300) at 4°C overnight. Cells were washed with PBS three times and incubated with goat anti-mouse Alexa Fluor 488 (1:500) at RT for 1 h. After another three rounds of washing, cover glasses were incubated with Vectashield Antifade Mounting medium with DAPI (Vector Laboratories), placed in the slides, and fixed using nail polish. Then, images displaying nuclear SREBP-1 and SREBP-2 were acquired using a Leica DMi8 microscope.

### Hepatic immunochemistry

ORO and HE staining in mouse OCT/paraffin-embedded liver tissue were carried out using 8 μm thick frozen sections and 4 μm paraffin sections. After placed at RT for 30 min, frozen sections were immersed in propylene glycol for 2 min and stained with pre-warmed ORO solution for 10 min. Following the staining procedure, tissues were differentiated in 85% propylene glycol for 1 min and then counterstained using modified Mayer’s hematoxylin and mounted using an aqueous mounting medium (Carl Roth). Given that lipids are dissolved by organic solvents during this sample preparation, ORO staining cannot be performed in paraffin sections. Hepatic HE staining in either paraffin or frozen sections was performed as described above for HE staining of vessels.

### Expression and purification of MIF and MIF-2 proteins

MIF was expressed in E. coli BL21/DE3-pET11b and purified essentially as described previously (56, 79). MIF-2 was cloned into a pET22b expression vector, expressed in *E. coli* BL21-CodonPlus, and purified following a slight modification of a previously reported procedure (30). Briefly, bacterial extracts were filtered, purified by Q Sepharose chromatography (Cytiva) applying fast protein liquid chromatography (Äkta-FPLC, Cytiva) and a subsequent high performance liquid chromatography step using C18 reverse-phase separation (RP-HPLC). MIF-2 was refolded based on the protocol established for MIF and following a previously described method (30). SDS-PAGE in combination with Coomassie or silver staining was utilized to evaluate protein purity and mass spectroscopy applied to verify protein identity. The resulting proteins contained <10 pg LPS/μg protein, as quantified by the PyroGene Recombinant Factor C assay (Cambrex).

### Flow cytometry

Whole mouse blood was collected by 30G needles into EDTA tubes, placed at RT for 10 min before storage on ice. Blood cells and plasma were separated by centrifugation at 300 g for 12 min at 4°C. To estimate percentages of different types of white blood cells (WBC), RBC lysis buffer at RT was used to lyse RBCs for 3 min, and filtered PBS with 0.5% BSA (i.e. FACS buffer) was used to wash and suspend WBC pellets. Then, an antibody cocktail panel (1:100) containing V450-conjugated anti-CD45 (BD Biosciences), FITC-conjugated anti-CD3 (Miltenyi Biotec), APC-Cy7-conjugated anti-CD19 (BioLegend), PE-Cy7-conjugated anti-CD11b (BioLegend), PE-conjugated anti-CD11c (BioLegend), APC-conjugated anti-Ly6C (BioLegend), and PerCP-conjugated anti-Ly6G (BioLegend), was applied to stain leukocyte subsets. After 45 min of antibody incubation, these stained cells were sorted in a FACSVerse™ (BD Biosciences) flow cytometer. Compensations and special gating strategies were set using isotype controls and fluorescence minus one (FMO) antibody combinations. Raw data were obtained by the FACSiva software (BD Biosciences), and analysis carried out via FlowJo software (Treestar).

### Plasma lipid and lipoprotein analysis

Concentrations of total triglycerides and total cholesterols in mouse plasma from different cohorts were enzymatically determined via triglyceride colorimetric assay kit (1:2) and cholesterol fluorometric assay kit (1:2000) following the manufacturer’s guidance. Proper dilution ratios of plasma are critical factors and were taken into account in both assays.

To analyze lipoprotein profiles including very low-density lipoprotein (VLDL), low-density lipoprotein (LDL), and high-density lipoprotein (HDL), plasma samples were subjected to FPLC chromatography using a Superose 6 size exclusion column (SEC, Cytiva). Different lipoprotein fractions were separated and collected according to their retention times as follows: VLDL between 40 and 50 min, LDL between 50 and 70 min, and HDL between 70 and 90 min. Separated lipoproteins were quantitated and results are presented as optical density (OD) measurements analyzing the absorbance at 492 nm using a plate reader (Perkin Elmer Enspire).

### Monocyte adhesion on aortic endothelial cell monolayers under flow conditions

Human aortic endothelial cells (HAoECs) were seeded at a density of 60,000 cells in collagen-precoated Ibidi µ-Slides I 0.8 and cultured overnight in order to obtain confluent monolayers and stimulated with 16 nM of human MIF or different concentrations of human MIF-2 (i.e. 0 nM, 0.8 nM, 1.6 nM, 4 nM, 8 nM, 16 nM, 40 nM and 80 nM) for 2 h, essentially as described previously (15, 56). In parallel, MonoMac6 cells (∼0.5 × 10^6^ cells/mL) were treated with the same concentration(s) of MIF or MIF-2 for 2 min. Then they were transferred into endothelial cell growth medium containing 10 mM HEPES and 0.5% BSA, and kept at 37°C. Before endothelial superfusion, MonoMac6 suspensions were adjusted to a final concentration of 1 mM HEPES by adding MgSO_4_ and CaCl_2_ solutions. Capitalizing on the Ibidi pump system and perfusion set, MonoMac6 suspensions were then used to perfuse corresponding flow channels containing HAoEC monolayers at a shear rate of 1.5 dyn/cm^2^ for 10 min at 37°C. Lastly, images from four different positions in the middle of each channel were acquired using a Leica DMi8 fluorescence microscope and adherent monocytes were quantified with Image J.

### Transwell migration assay

The chemotactic migration of primary murine B lymphocytes or human monocytes was assessed using a Transwell device as published previously (16). Human monocytes or splenic B cells were isolated as described above and maintained in RPMI 1640 medium with 10% FCS and 1% P/S overnight. A suspension of 100 µL containing 1 x 10^6^ cells was loaded in the upper chamber of the Transwell culture insert. Filters were transferred into the wells (‘the lower chamber’) containing different doses of MIF-2 with or without inhibitors. The inhibitory effects on MIF-2-mediated cell migration were determined by prior incubation of inhibitors at 37°C for 45 min. The Transwell device was incubated for 4 h at 37°C in a humidified atmosphere of 5% CO_2_. Cells migrated into the lower chamber were collected and counted by flow cytometry using the CountBright™ Absolute Counting Bead (Molecular Probes) method.

### 3D chemotaxis assay

The 3D migration of isolated human monocytes and primary B cells in response to MIF-2 (or MIF) was assessed using the 3D chemotaxis µ-Slide system (Ibidi GmbH) according to the manufactureŕs instructions. Freshly isolated cells (4 x 10^6^) were seeded in a pre-prepared gel matrix containing 1 mg/mL rat tail collagen type I (Merck Millipore) in DMEM. The collagen gel containing the cell suspension was incubated at 37°C for 30 min for polymerization and was subsequently subjected to a gradient of MIF-2 (0 nM, 2 nM, 4 nM, 8 nM, 16 nM and 32 nM) to evaluate the migratory behavior of monocytes to MIF-2.

For competition experiments comparing the potency of MIF-2 with that of MIF, splenic mouse B cells were used and competition between 4 nM MIF-2 and 8 nM MIF studied. To monitor cell motility in live mode, time-lapse imaging was performed every 1 min at 37°C for 2 h applying a Leica DMi8 fluorescence microscope. Original images were imported as stacks to ImageJ software and were analyzed through the manual tracking and the chemotaxis and migration tools (Ibidi GmbH).

### In vivo B-lymphocyte homing assay

Primary B lymphocytes were obtained from wild type (WT) mouse spleen as described above. 4-20 x 10^6^ cells per mouse were injected into non-irradiated 4- to 8-week-old *Apoe^−/−^* or *Mif^−/−^* mice through the tail vein. In the experimental group, B cells were treated with 4-IPP for 30 min before injection. For controls, B cells were incubated with a DMSO-containing control buffer. One hour after injection, mice were sacrificed, and blood, bone marrow, spleen, and lymph nodes (mesenteric, inguinal, brachial, axillary, and cervical) were collected. Injected B cells from different organs were quantitated by flow cytometry using human-specific anti-CD45 (1:100) and anti-CD19 (1:100) antibodies. Data was analyzed by FlowJo software and the homing effect was expressed by normalizing the number of B cells detected in 10^6^ cells. Non-injected mice were used as controls.

### Sequence alignment

MIF and MIF-2 sequence alignment was performed by the ClustalW algorithm (http://www.genome.jp/tools-bin/clustalw) using standard parameters in the Jalview multiple sequence alignment editor desktop application.

### CXCR4 internalization assay

Primary splenic B cells from mouse expressing CXCR4 on its surface were used in this assay. B lymphocytes from *Cd74^−/−^* mice were resuspended in RPMI1640 medium supplemented with 10% FCS and 1% P/S under a 5% CO_2_ humidified atmosphere at 37°C. These isolated cells were seeded at a density of 1x 10^6^/mL in 24-well plate. After stimulating with different dose of MIF-2 (i.e. 0 nM, 2 nM, 4 nM, 10 nM, 20 nM) and 4 nM MIF-2 at different time points (i.e. 0 min, 5 min, 10 min, 15 min, 20 min, 30 min), surface expressions of CXCR4 in different groups were measured by flow cytometry, through using isotype/fluorochrome matched controls meanwhile. The median fluorescence intensity (MFI) was quantified and compared through FlowJo software.

### CXCR4-specific yeast reporter signaling assay

The *S. cerevisiae* strain (CY12946) expressing functional human CXCR4 protein which replaces the yeast STE2 receptor and is linked to a MAP kinase signaling pathway and β-galactosidase (lacZ) read-out is a well-established human CXCR4-specific receptor binding and signaling cell system (55, 56). MIF was previously demonstrated to elicit a CXCR4-specific signaling in this system (55). In short, yeast transformants stably expressing human CXCR4 were grown at 30°C overnight in yeast nitrogen base selective medium (Formedium). Cells were diluted to an OD_600_ of 0.2 in yeast extract-peptone-dextrose (YPD) medium and grown until exponential phase with an OD_600_ of ∼0.3-0.6. Transformants were incubated with different concentrations of human MIF-2 (i.e. 0 µM, 2 µM, 4 µM, 8 µM, 16 µM, 32 µM) for 1.5 h. Moreover, 10 µM MIF-2 and 20 µM MIF were used to directly compare the binding of these proteins to CXCR4. β-galactosidase activity was measured by a commercial BetaGlo Kit (Promega) to assess the activation of CXCR4 signaling, and OD_600_ was detected by a plate reader with a 600 nm filter.

### Fluorescence spectroscopy

Fluorescence spectroscopic titrations of the soluble CXCR4 surrogate molecule msR4M-L1 with MIF-2 were performed using a JASCO FP-6500 fluorescence spectrophotometer. MIF-2 was used in aqueous buffer containing 20 mM sodium phosphate, pH 7.2. The CXCR4-derived peptide msR4M-L1 was freshly dissolved in 1,1,1,3,3,3-hexafluoro-2-propanol (HFIP) (Sigma-Aldrich) at 4°C at a concentration of 1 or 2 mM. For the titrations, the solutions consisted of Fluos-labeled peptide (5 nM) and different concentrations of MIF-2 in 10 mM sodium phosphate buffer (pH 7.4) containing 1% HFIP. The 500-600 nm spectra shown are representative of one of three independent experiments. App. K_D_ values were calculated assuming a 1:1 binding model (80) using sigmoidal curve fitting with OriginPro 2016 (OriginLab Corporation).

### DiI-LDL uptake assay in PBMC-derived macrophages

The uptake assay of DiI complex-labeled low-density lipoprotein (DiI-LDL) in primary human monocyte-derived macrophages was performed as previously described (56, 57). Briefly, macrophages were cultured in the full RPMI 1640 medium at 37°C, and then changed to MEM medium containing 0.2% BSA for 2-4 h to achieve starvation. Later, cells were pre-incubated with inhibitors (10 µM AMD3100, 4-IPP, or 4-CPPC) for 30 min along with 1 µg/mL MIF-2 overnight. The second day, cells were cultured in the same medium added with 1% 2-hydroxypropyl-β-cyclodextrin (HPCD) (Sigma-Aldrich) at 37°C for 45 min. After rinsing with PBS three times, macrophages were maintained in 25 µg/mL DiI-LDL solution at 4°C for 30 min (the ‘binding step’), and subsequently moved to 37°C for 20 min (the ‘uptake step’). Treated macrophages were washed with PBS, fixed with 4% PFA, permeabilized with 0.1% Triton X in PBS, and stained with DAPI (1:100,000). Representative images were acquired with a Leica DMi8 fluorescence microscope and the uptake effect was characterized as the index of relative corrected total cell fluorescence (CTCF) via ImageJ and Excel.

### Native LDL uptake assay in Huh-7 hepatocytes

1 x 10^4^ Huh-7 cells were seeded in 24-well plates on the day before starting the experiment and then starved with 2% FCS DMEM overnight. On day 3, cells were stimulated with 8 nM MIF-2 with or without IgG, AMD3100 and LN-2, and cultured in 2% FCS DMEM for 24 h. After treatment, cells were incubated with 10 µg/mL human LDL at 37°C for 4 h, followed by 3 times of washing in PBS. Next, cells were fixed in 4% PFA and subsequently stained with ORO solution for 5 min. Stained cells were washed and counterstained with DAPI (1:100,000) and mounted on a glass slide. Images were captured using a Leica DMi8 microscope.

### Human carotid artery plaques

Human carotid artery plaques were retrieved during carotid artery endarterectomy (CEA) surgery at the Department of Vascular and Endovascular Surgery at the Klinikum rechts der Isar of the Technical University Munich (TUM). Advanced carotid lesions with an unstable/ruptured or stable plaque phenotype (determined as previously described (81, 82) were cut in ∼50mg pieces on dry ice. All patients provided their written informed consent. The study was approved by the local ethics committee at the Medical Faculty of the Klinikum rechts der Isar of the Technical University Munich (ethics approval 2799-10).

### Statistics

Statistical analysis was performed using GraphPad Prism version 8 software. Data are represented as means ± SD. After testing for normality, data were analyzed by two-tailed Student’s t-test, Mann-Whitney U, or Kruskal-Wallis test as appropriate. Differences with *P* < 0.05 were considered to be statistically significant.

## Supplemental Information

### 1. SUPPLEMENTAL METHODS

Reagent and Resources Table.

**Reagent and Resources Table.**
| REAGENT or RESOURCE | SOURCE | IDENTIFIER |
| --- | --- | --- |
| Antibodies |  |  |
| Anti-mouse CD45 V450 (clone 30-F11) | BD Biosciences | Cat# 560501 |
| Anti-mouse CD3 FITC (clone REA641) | Miltenyi Biotec | Cat# 130-119-758 |
| Anti-mouse CD19 APC/Cy7 (clone 6D5) | BioLegend | Cat# 115530 |
| Anti-mouse CD11c PE (clone N418) | BioLegend | Cat# 117308 |
| Anti-mouse/human CD11b PE/Cy7 (clone M1/70) | BioLegend | Cat# 101216 |
| Anti-mouse Ly-6C APC (clone HK1.4) | BioLegend | Cat# 128016 |
| Anti-mouse Ly-6G PerCP (clone 1A8) | BioLegend | Cat# 127654 |
| Anti-SREBP-1 mouse monoclonal antibody | Santa Cruz Biotechnology | Cat# sc-17755 |
| Anti-SREBP-2 mouse monoclonal antibody | BD Biosciences | Cat# 557037 |
| Anti-FASN mouse monoclonal antibody | Santa Cruz Biotechnology | Cat# sc-55580 |
| Anti-LDLR mouse monoclonal antibody | Santa Cruz Biotechnology | Cat# sc-18823 |
| Anti-CD68 rat monoclonal antibody | Bio-Rad Laboratories | Cat# MCA1957GA |
| Anti-AMPKα rabbit monoclonal antibody | Cell Signaling Technology | Cat# 5832S |
| Anti-phospho-AMPKα rabbit monoclonal antibody | Cell Signaling Technology | Cat# 2535 |
| Anti-β-actin mouse monoclonal antibody | Santa Cruz Biotechnology | Cat# sc-47778 |
| Anti-CXCR4 rabbit polyclonal antibody | Invitrogen | Cat# PA3-305 |
| Anti-CD74 mouse monoclonal antibody | Santa Cruz Biotechnology | Cat# sc-6262 |
| Anti-D-DT rabbit polyclonal antibody | Bucala lab (Yale) | Not applicable |
| Alexa Fluor 647-conjugated donkey anti-rat IgG (H+L) | Jackson ImmunoResearch Europe Ltd. | Cat# 712-606-153 |
| Alexa Fluor 488-conjugated goat anti-rat IgG (H+L) | Invitrogen | Cat# A-11006 |
| Alexa Fluor 555-conjugated goat anti-rabbit IgG (H+L) | Invitrogen | Cat# A-21429 |
| Goat anti-mouse HRP-conjugated antibody | Abcam | Cat# ab6789 |
| Goat anti-rabbit HRP-conjugated antibody | GE Healthcare | Cat# NA934V |
| Chemicals and inhibitors |  |  |
| 4-CPPC (MIF-2-selective inhibitor) | Aurora Fine Chemicals LLC | Cat# K00.223.811<br>29553-70-6 |
| 4-IPP (non-selective MIF/MIF-2 inhibitor) | Tocris Bioscience | Cat# 3429 |
| Low density lipoprotein from human plasma, Dil complex (Dil LDL) | Thermo Fisher Scientific | Cat# L3482 |
| Low density lipoprotein from human plasma (LDL) | Thermo Fisher Scientific | Cat# L3486 |
| Paraformaldehyd (PFA) 4% in PBS 7.4 | Morphisto GmbH | Cat# 1176201000 |
| TRIzol™ reagent | Thermo Fisher Scientific | Cat# 15596018 |
| CountBright™ Absolute Counting Beads, for flow cytometry | Thermo Fisher Scientific | Cat# C36950 |
| PolyFect transfection reagent | Qiagen | Cat# 301107 |
| Recombinant human M-CSF | PeproTech GmbH | Cat# 300-25 |
| RBC lysis buffer (10X) | BioLegend | Cat# 420301 |
| Oil Red O solution 0.5% in propylene glycol | Sigma-Aldrich | Cat# O1516-500ML |
| Ficoll Paque Plus | GE Healthcare | Cat# GE17-1440-02 |
| Albumin fraction V | Carl Roth | Cat# 8076.4 |
| Methanol | Carl Roth | Cat# 0082.2 |
| CozyHi™ prestained protein ladder | HighQu GmbH | Cat# PRL0202 |
| TWEEN 20 viscous liquid | Sigma-Aldrich | Cat# P2287-500ML |
| Donkey serum | Sigma-Aldrich | Cat# D9663-10ML |
| 30% Bis acrylamide solution | Bio-Rad Laboratories | Cat# 1610156 |
| Hematoxylin solution, Mayer's | Sigma-Aldrich | Cat# MHS80-2.5L |
| Eosin Y solution, aqueous | Sigma-Aldrich | Cat# HT110232-1L |
| DL-dithiothreitol ≥98% (HPLC), ≥99.0% | Sigma-Aldrich | Cat# D0632-25G |
| 2-Propanol ROTISOLV® ≥99.95 %, LC-MS-Grade | Carl Roth | Cat# AE73.2 |
| NuPAGE transfer buffer (20X) | Thermo Fisher Scientific | Cat# NP00061 |
| Xylol (Isomergemisch) | VWR International | Cat# 28976.294 |
| Water nuclease-free | VWR International | Cat# 436912C |
| Bouin's solution | Sigma-Aldrich | Cat# HT10132-1L |
| Sulphuric acid | Carl Roth | Cat# KK74.1 |
| TEMED | Bio-Rad Laboratories | Cat# 1610801 |
| Penicillin-streptomycin (5,000 U/mL) | Invitrogen | Cat# 15140-122 |
| Fetal bovine serum | Thermo Fisher Scientific | Cat# 10091148 |
| Pierce protease inhibition tablets | Thermo Fisher Scientific | Cat# A32963 |
| Medium (cell culture) |  |  |
| Dulbecco's phosphate buffered saline modified, without calcium chloride and magnesium chloride | Sigma-Aldrich | Cat# D8537 |
| Dulbecco's phosphate-buffered saline | Sigma-Aldrich | Cat# D8662 |
| RPMI 1640 Medium GlutaMax + supplements | Invitrogen | Cat# 61870-044 |
| DMEM/F-12, GlutaMAX™ supplement | Invitrogen | Cat# 31331-093 |
| Mounting medium (staining) |  |  |
| Fluoromount™ aqueous mounting medium | Sigma-Aldrich | Cat# F4680-25ML |
| Kaiser's glycerol gelatine phenol-free | Carl Roth | Cat# 6474.1 |
| Eukitt® Quick-hardening mounting medium | Sigma-Aldrich | Cat# 25608-33-7 |
| Vectashield Antifade mounting medium with DAPI | Vector Laboratories | Cat# H-1200 |
| Critical commercial kits |  |  |
| Proteome Profiler Mouse Cytokine Array Kit, panel A | R&D systems | Cat# ARY006 |
| First strand cDNA synthesis Kit | Thermo Fisher Scientific | Cat# K1612 |
| ORA™ SEE qPCR Green ROX L Mix, 2X | HighQu GmbH | Cat# QPD0150 |
| Triglyceride colorimetric assay kit | Cayman Chemical | Cat# 10010303 |
| Cholesterol fluorometric assay kit | Cayman Chemical | Cat# 10007640 |
| Trichrome stain (Masson) kit | Sigma-Aldrich | Cat# HT15-1KT |
| RNA/protein purification Plus Kit | Norgen Biotek Corp. | Cat# 48200 |
| Pan B cell Isolation kit, mouse | Miltenyi Biotec | Cat# 130095813 |
| Pan monocyte isolation kit, human | Miltenyi Biotec | Cat# 130096537 |
| Pierce™ TMB substrate kit | Thermo Fisher Scientific | Cat# 34021 |
| SuperSignal West Femto Maximum Sensitivity substrate | Thermo Fisher Scientific | Cat# 34095 |
| Weigert's iron hematoxylin set | Sigma-Aldrich | Cat# 537864 |
| SuperSignal West Dura Extended<br>Duration substrate | Thermo Fisher Scientific | Cat# 34076 |
| Experimental models |  |  |
| C57BL/6J mice | The Jackson Laboratory | Not applicable |
| *ApoE*<sup>-/-</sup> mice on a C57BL/6 background | The Jackson Laboratory | Not applicable |
| *Mif-2*<sup>-/-</sup> *ApoE*<sup>-/-</sup> mice on a C57BL/6 background | This paper | Not applicable |
| *Mif*<sup>-/-</sup> mice on a C57BL/6 background | This paper | Not applicable |
| Equipment |  |  |
| BD FACSVerse™ flow cytometer | BD Biosciences | Cat# 651155 |
| Mini gel tank | Invitrogen | Cat# A25977 |
| Mini blot module | Invitrogen | Cat# B1000 |
| Power supply EV3020 | Carl Roth | Cat# EV3020 |
| Odyssey Fc imaging system | LI-COR Biosciences | Cat# OFC-0976 |
| Heraeus™ Megafuge™ 16R centrifuge | Thermo Fisher Scientific | Cat# 75004270 |
| EnSpire Multimode plate reader | Perkin Elmer LAS GmbH | Cat# 23001395 |
| NanoDrop One Microvolume UV-Vis spectrophotometer | Thermo Fisher Scientific | Cat# AZY1602185 |
| Biometra TRIO 48 Touch thermocycler | Analytik jena GmbH | Cat# 2070723 |
| Horizontal gel electrophoresis system | Life Technologies | Cat# 41060 |
| Centrifuge 5424 R | Eppendorf AG | Cat# 5404000210 |
| Vortex shaker, VV3 | VWR International GmbH | Cat# 444-0007 |
| Thermal Shake Lite | VWR International GmbH | Cat# 460-0249P |
| Heracell™ VIOS 160i CO<sub>2</sub> incubator | Thermo Fisher Scientific | Cat# 51030285 |
| Biosafety cabinet | Kojair Tech Oy | Blue series |
| TC20™ automated cell counter | Bio-Rad Laboratories GmbH | Cat# 1450102 |
| Leica DMI8 fluorescent microscope | Leica Microsystems CMS GmbH | Cat# S/N 425074 |
| Zeiss Axio Imager M2 | Carl Zeiss AG | Cat# M2 |
| Confocal microscope | Carl Zeiss AG | Cat# LSM 880 |
| Leica TCS SP8 DIVE multiphoton microscope | Leica Microsystems CMS GmbH | Cat# Leica TCS SP8 |
| Cryotome Leica CM1950 platform | Leica Microsystems CMS GmbH | Cat# Leica CM1950 |
| Leica dissection microscope | Leica Microsystems CMS GmbH | Cat# Leica M125 C |
| Embedding Machine STP 120 | Leica Microsystems CMS GmbH | Cat# STP-120 |
| Rotor-Gene Q | Qiagen | Cat# R0915111 |
| TissueLyser LT | Qiagen | Cat# 23.1001/07538 |
| Software |  |  |
| GraphPad Prism 8 software | GraphPad | Version 8.4.3 |
| Rotor-Gene 6000 Series 1.7 software | Qiagen | Version 2.3.1 |
| BD FACS Diva software | BD Biosciences | Version 1.0.6.5230 |
| FlowJo V10 software | FlowJo | Version 17.0.2.0 |
| Image Studio™ software | LI-COR Biosciences | Version 5.2.5 |
| ImageJ software/Fiji software | NIH | https://imagej.nih.gov/ij/ |
| Leica application suite X software | Leica Microsystems CMS GmbH | Version 3.0.15878.1 |
| Carl Zeiss ZEN 2010 | Carl Zeiss AG | Version 2.3.64.0 |
| BioRender | BioRender | N/A |
| Other |  |  |
| Mouse diet: Western-type high fat diet | ssniff | Cat# E15721-347 |

### 2. SUPPLEMENTAL TABLES

**Supplemental Table S1.** Blood cell count, body weight and serum lipid levels from female *Apoe^−/−^* and *Mif-2^−/−^Apoe^−/−^* mice on cholesterol-rich high-fat diet (HFD) for 4.5 and 12 weeks. Shown are means ± SD. *P*-values calculated by Student’s t-test. Leukocytes were identified as CD45+; T lymphocytes as CD45+CD3+; B lymphocytes as CD45+CD19+; neutrophils as CD45+CD11b+Ly6G+; monocytes as CD45+CD11b+Ly6C+.

| Female, 4.5-week HFD | *Apoe*<sup>-/-</sup> | *Mif-2*<sup>-/-</sup>*Apoe*<sup>-/-</sup> | *P* value |
| --- | --- | --- | --- |
| **Serum lipid levels** |  |  |  |
| Triglycerides (mg/dL) | 188.135 ± 12.244 | 160.297 ± 36.357 | 0.0197 |
| Cholesterol (mg/dL) | 887.799 ± 104.492 | 703.902 ± 138.302 | 0.0013 |
| **Blood cell percentages (%)** |  |  |  |
| Lymphocytes (%) | 59.107 ± 1.929 | 63.339 ± 2.334 | 0.0313 |
| T lymphocytes (%) | 17.713 ± 4.350 | 26.363 ± 5.052 | 0.0409 |
| B-lymphocytes (%) | 41.387 ± 5.159 | 36.943 ± 4.546 | 0.2437 |
| Neutrophils (%) | 22.283 ± 4.952 | 24.212 ± 4.296 | 0.5777 |
| Monocytes (%) | 3.711 ± 1.131 | 1.493 ± 0.519 | 0.0118 |
| **Body weight** |  |  |  |
| Weight (g) | 24.420 ± 3.487 | 20.886 ± 1.254 | 0.0222 |
| Female, 12-week HFD | *Apoe*<sup>-/-</sup> | *Mif-2*<sup>-/-</sup>*Apoe*<sup>-/-</sup> | *P* value |
| **Serum lipid levels** |  |  |  |
| Triglycerides (mg/dL) | 189.344 ± 8.910 | 180.375 ± 7.116 | 0.0343 |
| Cholesterol (mg/dL) | 1002.305 ± 113.935 | 850.557 ± 92.842 | 0.0077 |
| **Blood cell percentages (%)** |  |  |  |
| Lymphocytes (%) | 69.461 ± 8.879 | 73.228 ± 7.893 | 0.3618 |
| T lymphocytes (%) | 24.519 ± 10.904 | 25.970 ± 3.704 | 0.7248 |
| B-lymphocytes (%) | 44.765 ± 5.474 | 47.168 ± 6.488 | 0.4061 |
| Neutrophils (%) | 13.189 ± 6.990 | 17.682 ± 6.487 | 0.1811 |
| Monocytes (%) | 3.482 ± 2.347 | 2.627 ± 0.700 | 0.3368 |
| **Body weight** |  |  |  |
| Weight (g) | 29.750 ± 2.914 | 25.775 ± 3.199 | 0.0140 |

#### Gating strategy was as follows

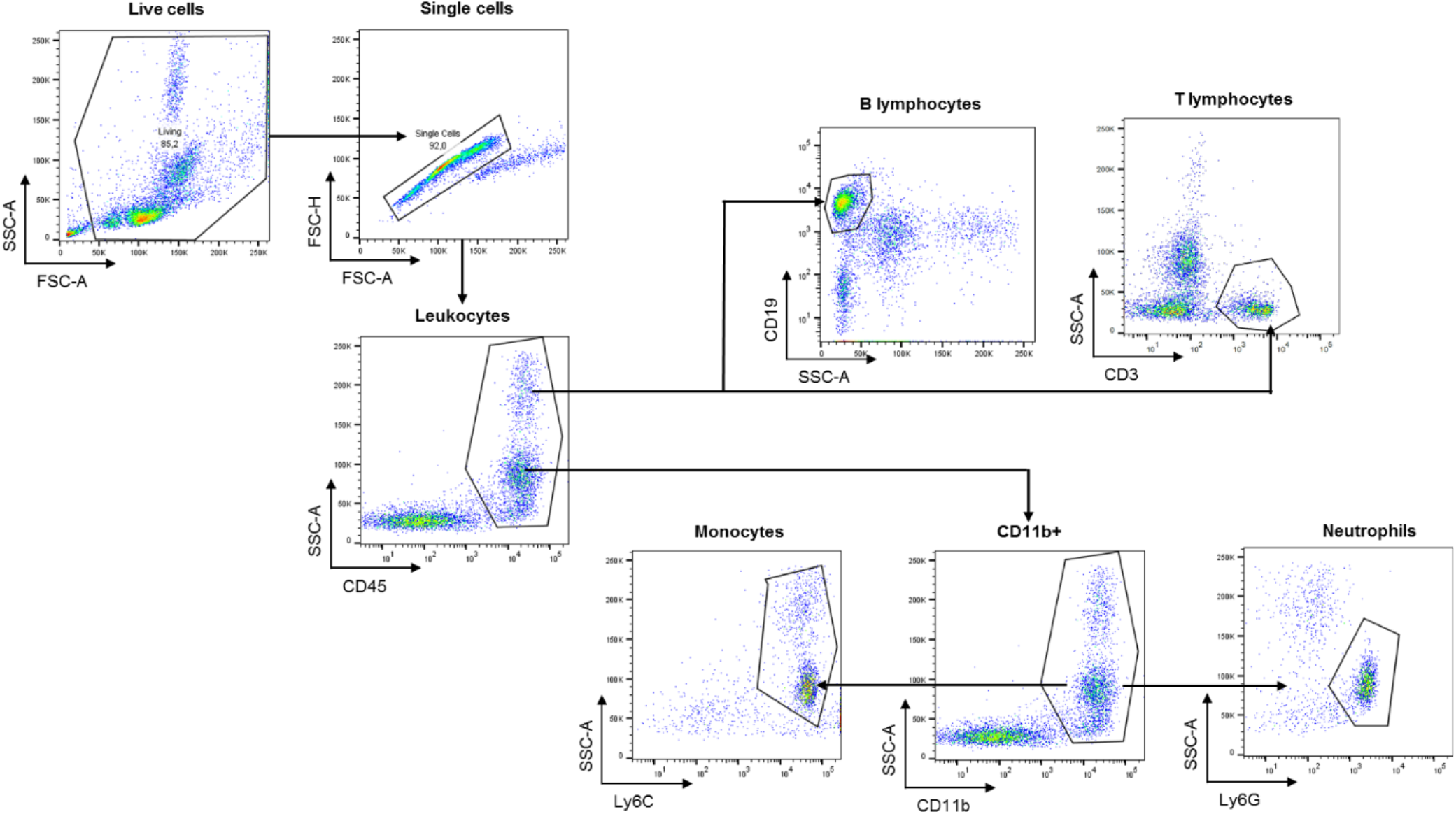

**Supplemental Table S2.**
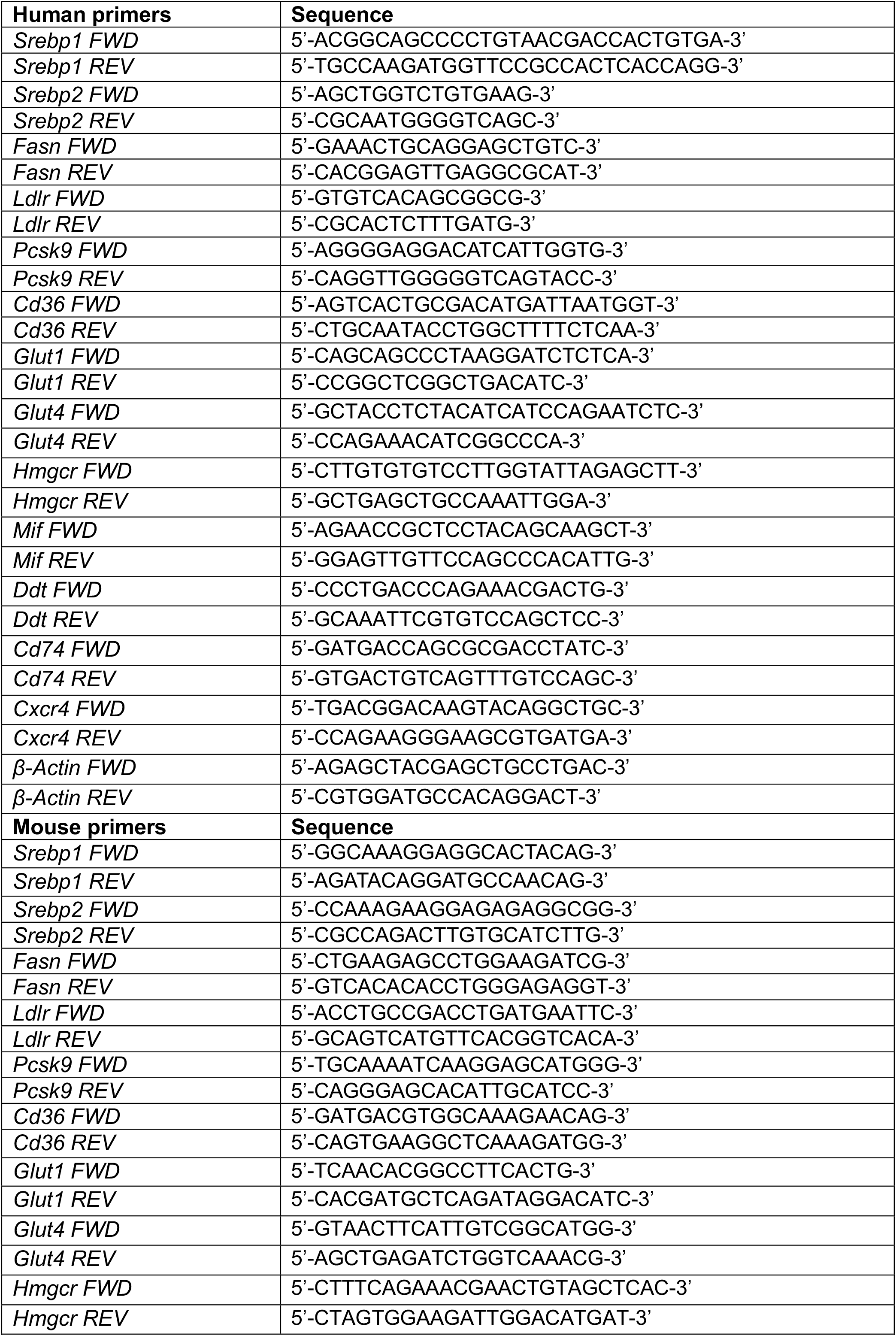
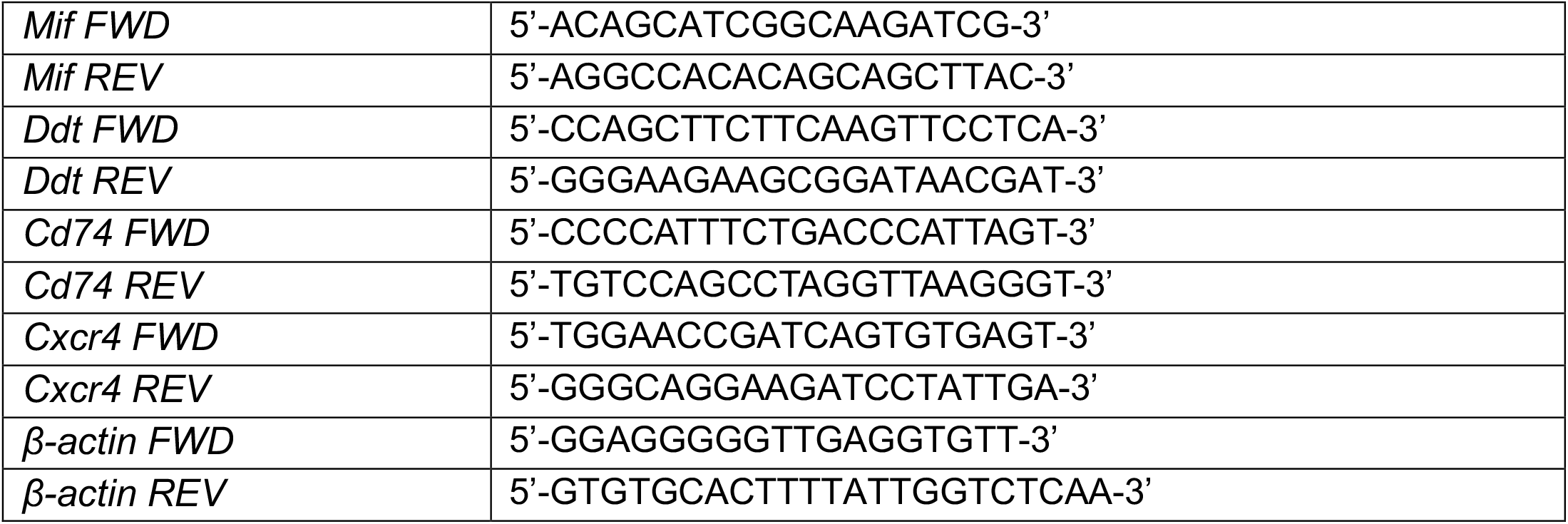
List of primers and sequences.

### 3. SUPPLEMENTAL FIGURES

**Figure S1.**
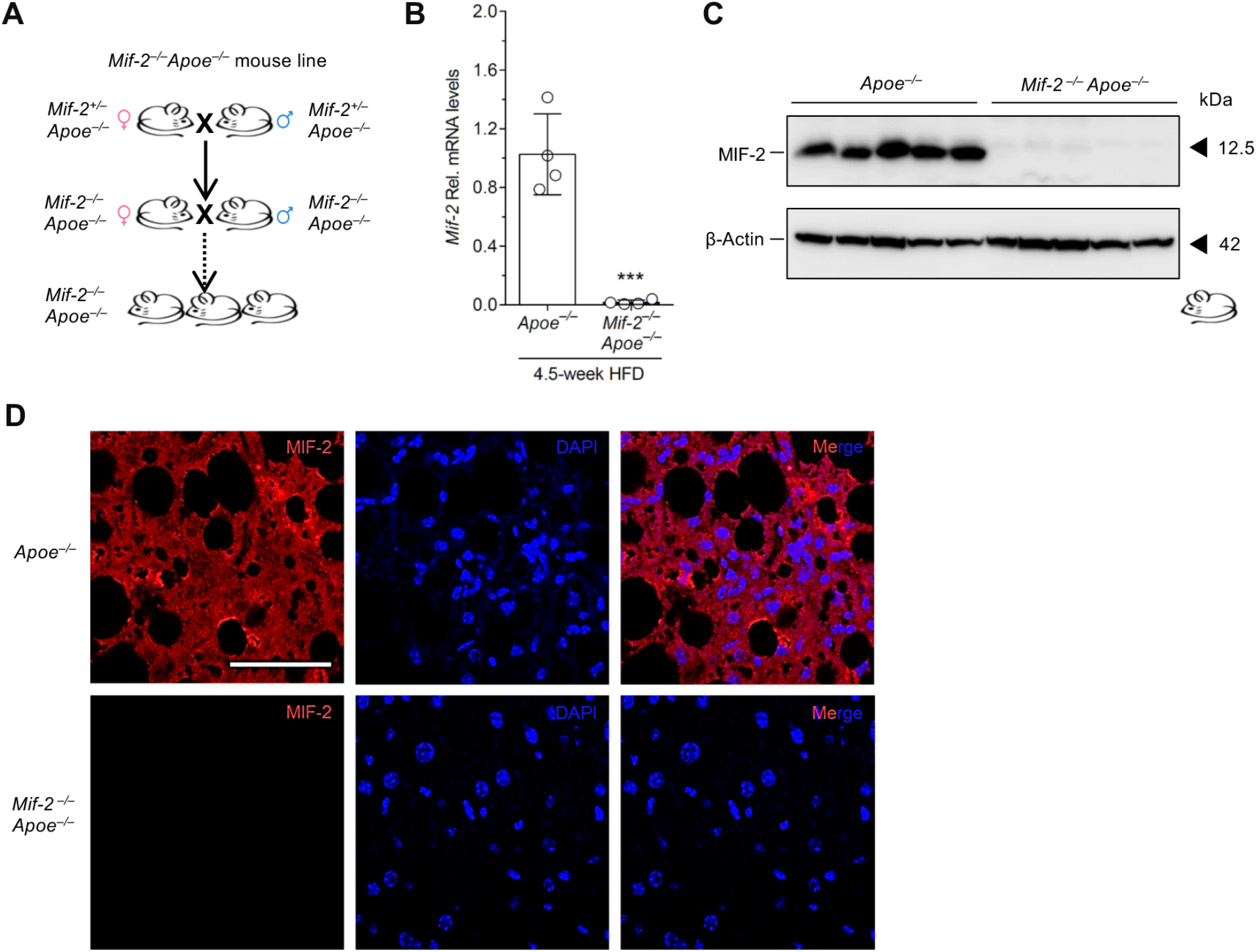
Generation and verification of the genetic *Mif-2* knockout in hyperlipidemic *Apoe*^−/−^ mice. (A) Schematics showing how the *Mif-2*^−/−^*Apoe*^−/−^ mouse line was bred and generated. (B) RT-qPCR-based genotyping on heart tissue from female *Apoe*^−/−^ and *Mif-2*^−/−^ *Apoe*^−/−^ mice to evaluate the global deletion of *Mif-2* on mRNA level. n = 4 mice per group. (C) Western blot was performed on liver tissue from female *Apoe*^−/−^ and *Mif-2*^−/−^*Apoe*^−/−^ mice to evaluate the global deletion of MIF-2 on protein level. n = 5 mice per group. (D) Fluorescent staining was performed on paraffin liver sections from female *Apoe*^−/−^ and *Mif-2*^−/−^*Apoe*^−/−^ mice after 12 weeks of Western-type cholesterol-rich high-fat diet (HFD) to visualize the global deletion of MIF-2 on cell/tissue level. Scale bar, 50 µm. All values are means ± SD; ***p<0.001.

**Figure S2.**
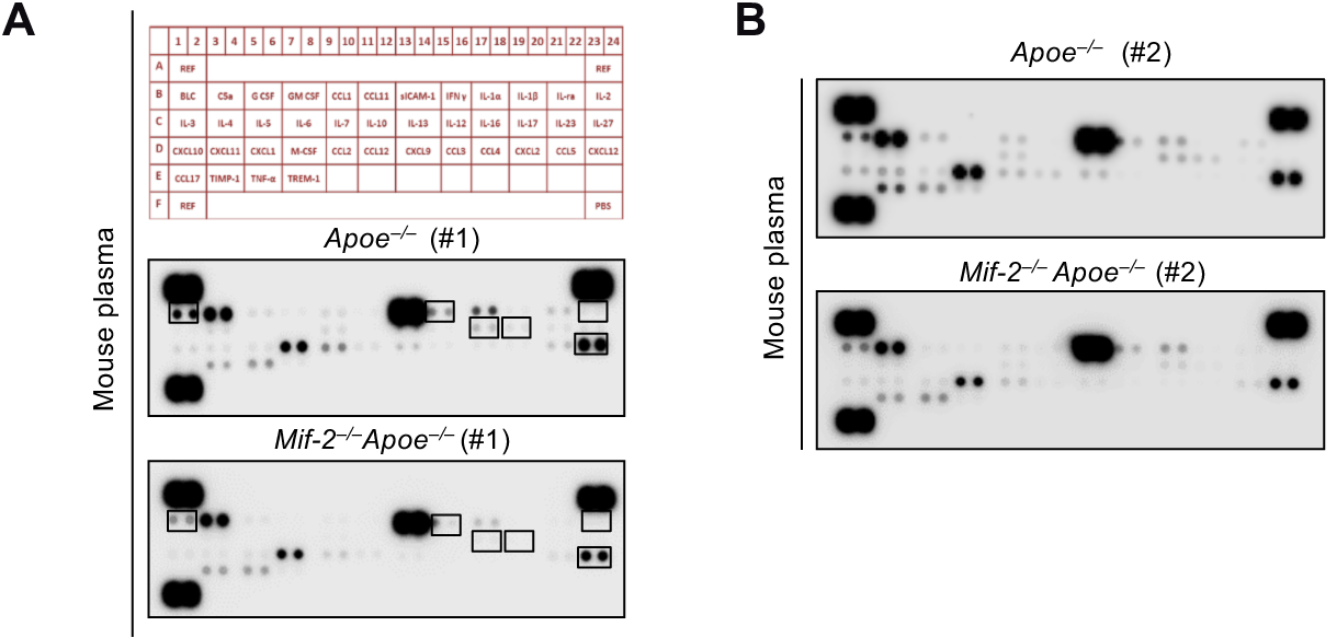
*Mif-2* knockout leads to downregulated inflammatory cytokine levels in female atherosclerotic mice. (A) Representative dot blots of differently expressed cytokines/chemokines from both female *Apoe*^−/−^ and *Mif-2*^−/−^*Apoe*^−/−^ mice fed a Western-type cholesterol-rich high-fat diet (HFD) for 4.5 weeks using a cytokine array approach. Two representative blots from each group are shown. Overall, plasma from n = 4 mice per group were analyzed.

**Figure S3.**
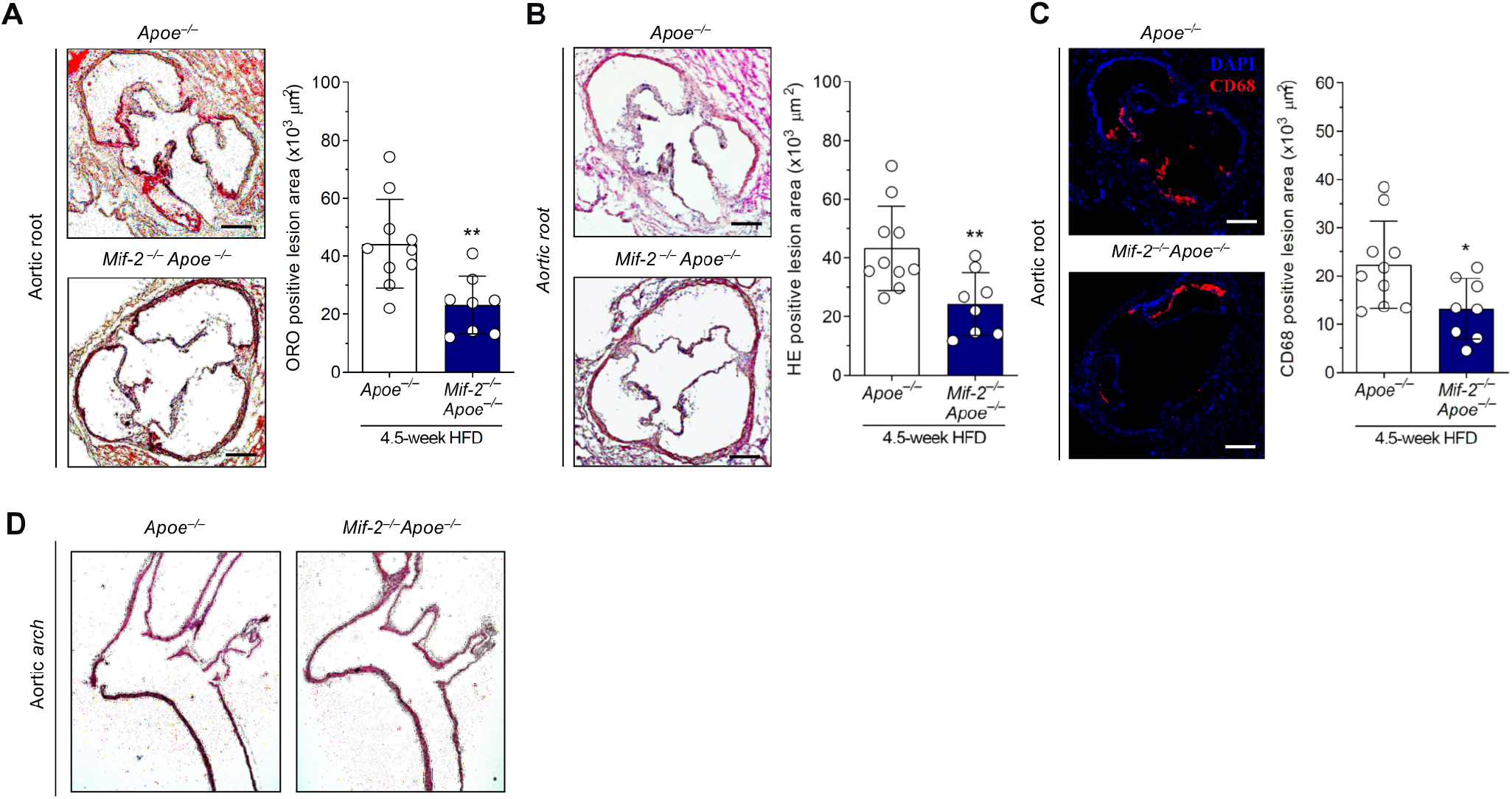
*Mif-2* deficiency attenuates early atherogenesis and vascular inflammation in male mice. (A) Representative Oil Red O (ORO) staining images of aortic roots from serial frozen sections (8 µm) of male *Apoe*^−/−^ and *Mif-2*^−/−^*Apoe*^−/−^ mice fed a Western-type cholesterol-rich high-fat diet (HFD) for 4.5 weeks and corresponding quantification results (12 sections per mouse). n = 8-10 mice per group. Scale bar, 250 µm. (B) Same as (A) except that hematoxylin-eosin (HE) staining was performed. (C) Representative images and quantification results of CD68^+^ macrophage content (red) in aortic root sections (8 µm) from male *Apoe*^−/−^ and *Mif-2*^−/−^ *Apoe*^−/−^ mice fed a Western-type cholesterol-rich high-fat diet (HFD) for 4.5 weeks (DAPI, blue). n = 8-10 mice per group. Scale bar, 250 µm. (D) Representative HE-stained images of aortic arch in paraffin sections (4 µm) from male *Apoe*^−/−^ and *Mif-2*^−/−^*Apoe*^−/−^ mice fed a Western-type cholesterol-rich high-fat diet (HFD) for 4.5 weeks. n = 8-10 mice per group. Scale bar, 250 µm. All values are means ± SD; *p<0.05; **p<0.01.

**Figure S4.**
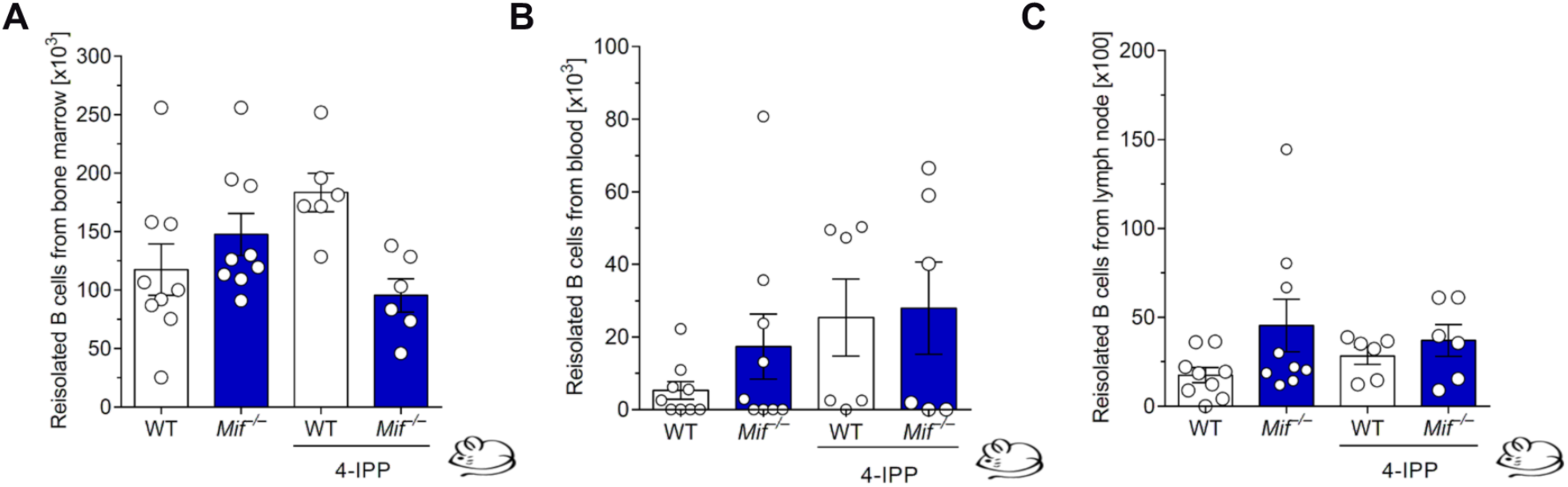
*In vivo* homing of B cells into bone marrow, blood, and lymph nodes. The experiment was performed according to the flow chart in Figure 2H. (A) Primary B lymphocyte homing into bone marrow. (B) Primary B lymphocyte in blood. (C) Primary B lymphocyte homing into lymph node. Groups of wildtype (WT) *versus Mif*^−/−^ and 4-IPP-*versus* vehicle-treated mice are compared.

**Figure S5.**
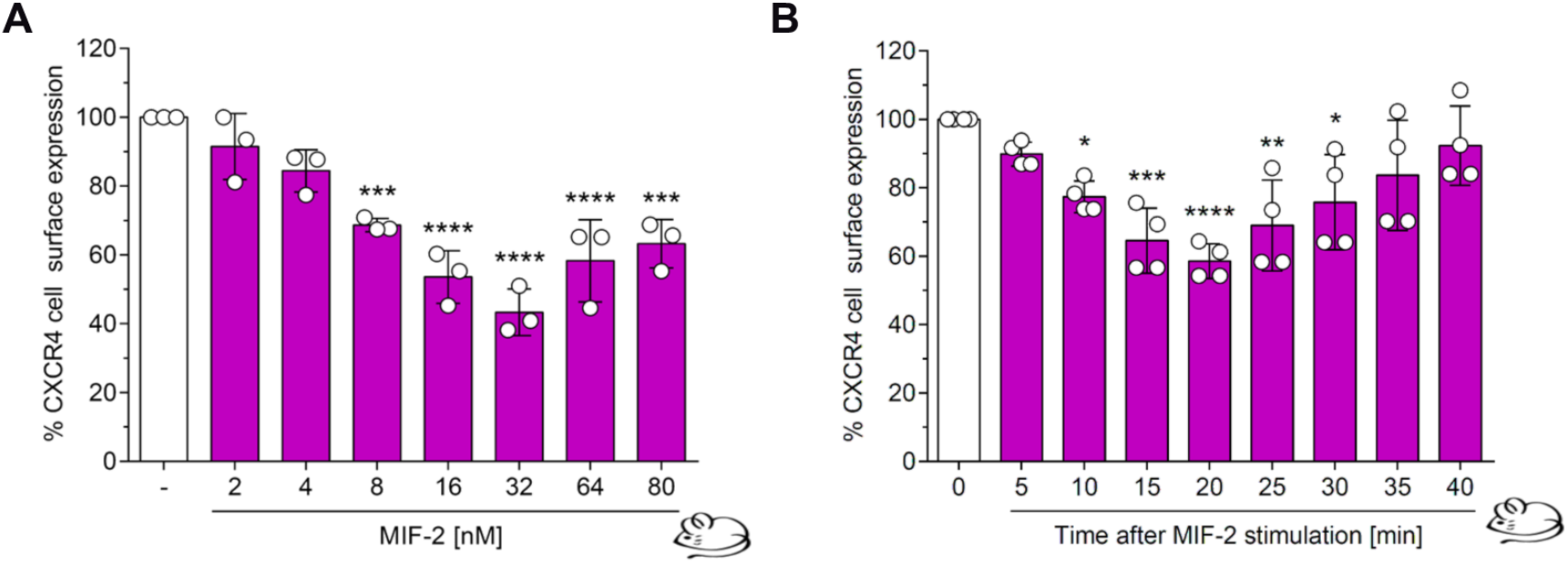
MIF induces CXCR4 internalization in a dose- and time-dependent manner. (A) MIF induces the internalization of CXCR4 in primary B lymphocytes from wildtype C57BL/6 mice in a dose-dependent manner. Concentration-dependency of MIF as indicated. (B) Same as (A) except that a time course experiment with 32 nM MIF was performed. All values are means ± SD; *p<0.05; **p<0.01; ***p<0.001; ****p<0.0001.

**Figure S6.**
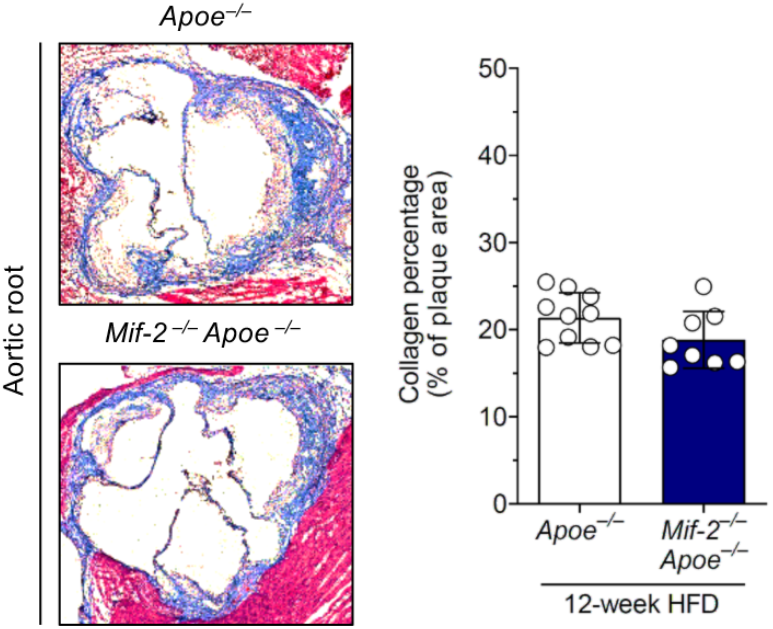
*Mif-2* deficiency does not affect the collagen content in hyperlipidemic *Apoe^−/−^* mice. (A) Representative images and quantification results of Masson-stained frozen sections in aortic root from *Apoe^−/−^* and *Mif-2^−/−^Apoe^−/−^* mice on 12-week Western-type cholesterol-rich high-fat diet (HFD). n=8-10 mice per group.

**Figure S7.**
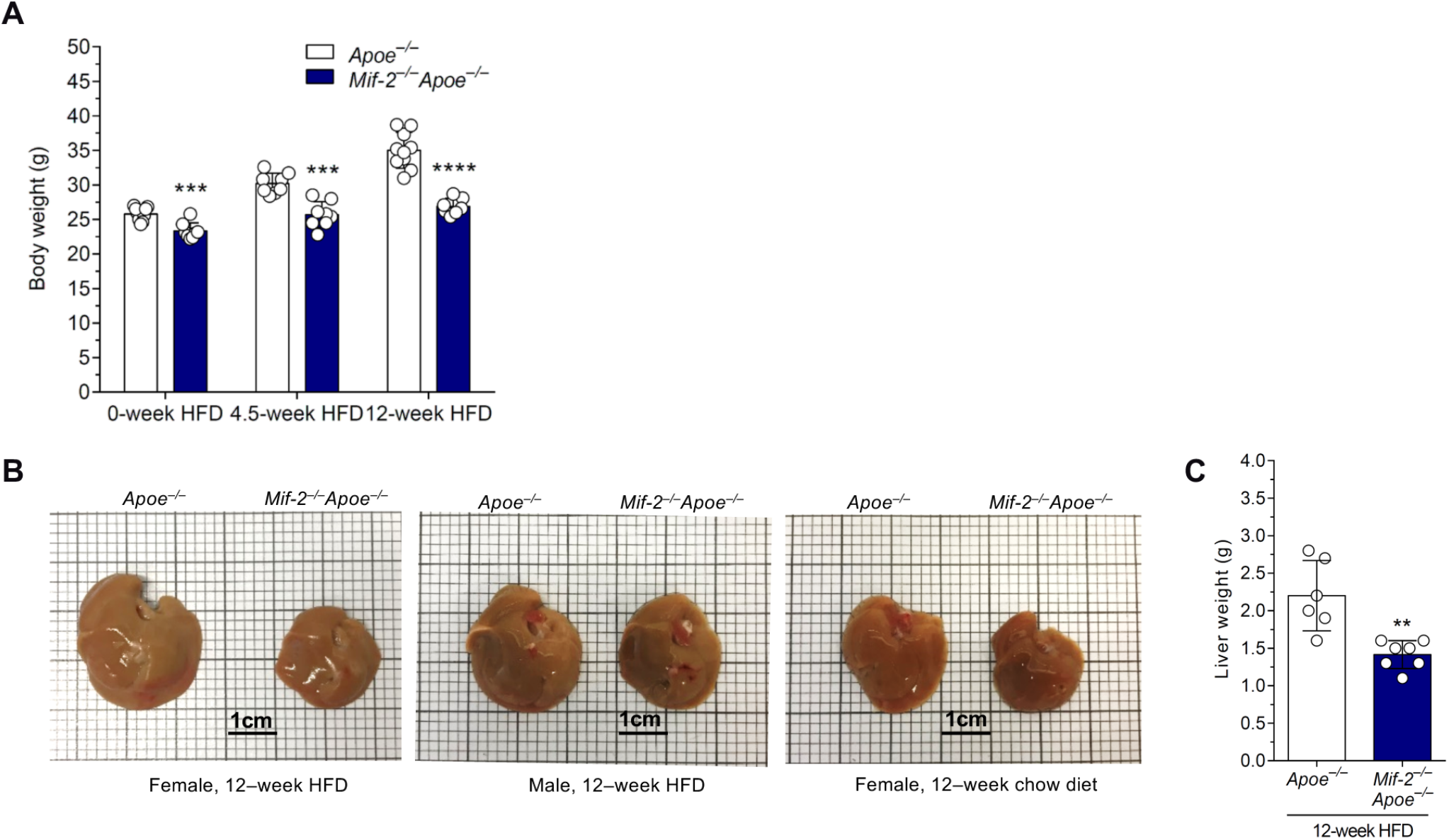
*Mif-2* knockout leads to weight loss and smaller liver size in male and female *Apoe^−/−^* mice. (A) Male *Apoe^−/−^* and *Mif-2^−/−^Apoe^−/−^* mice were fed Western-type cholesterol-rich high-fat diet (HFD) for 4.5 and 12 weeks. Significant differences in body weight are observed between both groups of male mice after 4.5- and 12-week HFD, as well as before onset of the HFD. (B) Representative images showing the livers of female and male *Apoe^−/−^* and *Mif-2^−/−^Apoe^−/−^* mice both after HFD and upon chow diet only. Livers of *Mif-2^−/−^Apoe^−/−^* mice appeared generally smaller. in both genders. Scale bar, 1 cm. (C) Quantification results showing that there is a significant decrease in liver weights in male *Mif-2^−/−^Apoe^−/−^* mice after 12-week HFD compared to *Apoe^−/−^* mice. All values are means ± SD; **p<0.01; ***p<0.001; ****p<0.0001.

**Figure S8.**
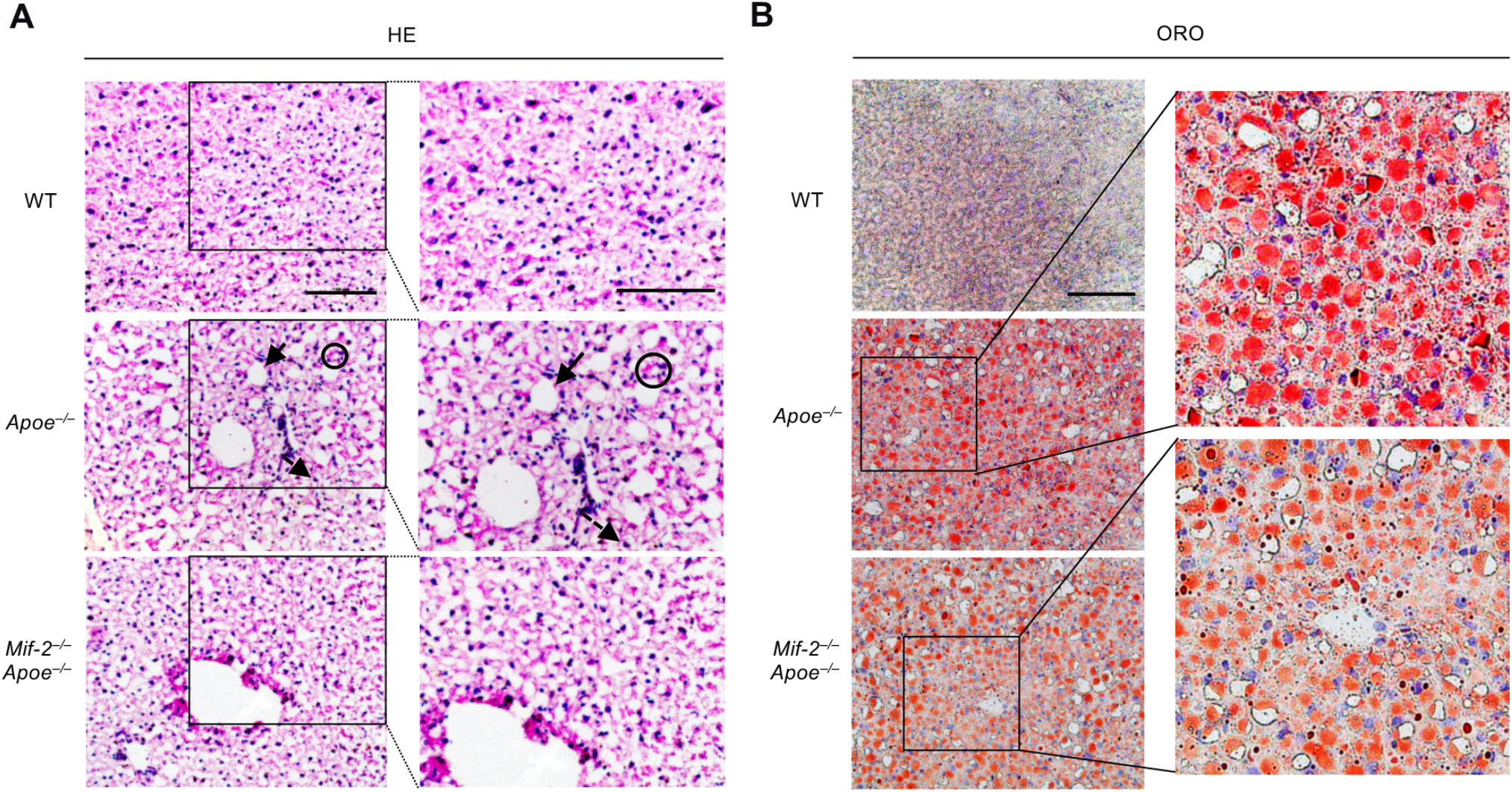
*Mif-2* knockout alleviates hepatic lipid accumulation in hyperlipidemic *Apoe^−/−^* mice. (A) Determination of hepatic lipid accumulation by HE staining using frozen sections. Representative liver images from *Mif-2^−/−^Apoe^−/−^* mice indicated reduced lipid accumulation compared with *Apoe^−/−^* mice. Classical structures for example macrovesicular steatosis (bold arrow), microvesicular steatosis (dotted line arrow), and clusters (aggregates) of inflammatory cells (within circles) could be identified. Right-hand panel, magnified images.(B) Analysis of hepatic lipid accumulation by ORO staining using frozen sections. Similar to HE staining, ORO-stained liver images from *Mif-2^−/−^Apoe^−/−^* mice exhibited reduced neutral lipid accumulation compared with *Apoe^−/−^* mice. Magnified insets on right-hand panel. The figure shows magnified views of the analysis shown in Figure 5C. Control images from wildtype (WT) mice (top left) without atherogenic background and Western-type high-fat diet (HFD) demonstrate the effect of the atherogenic phenotype on hepatosteatosis.

**Figure S9.**
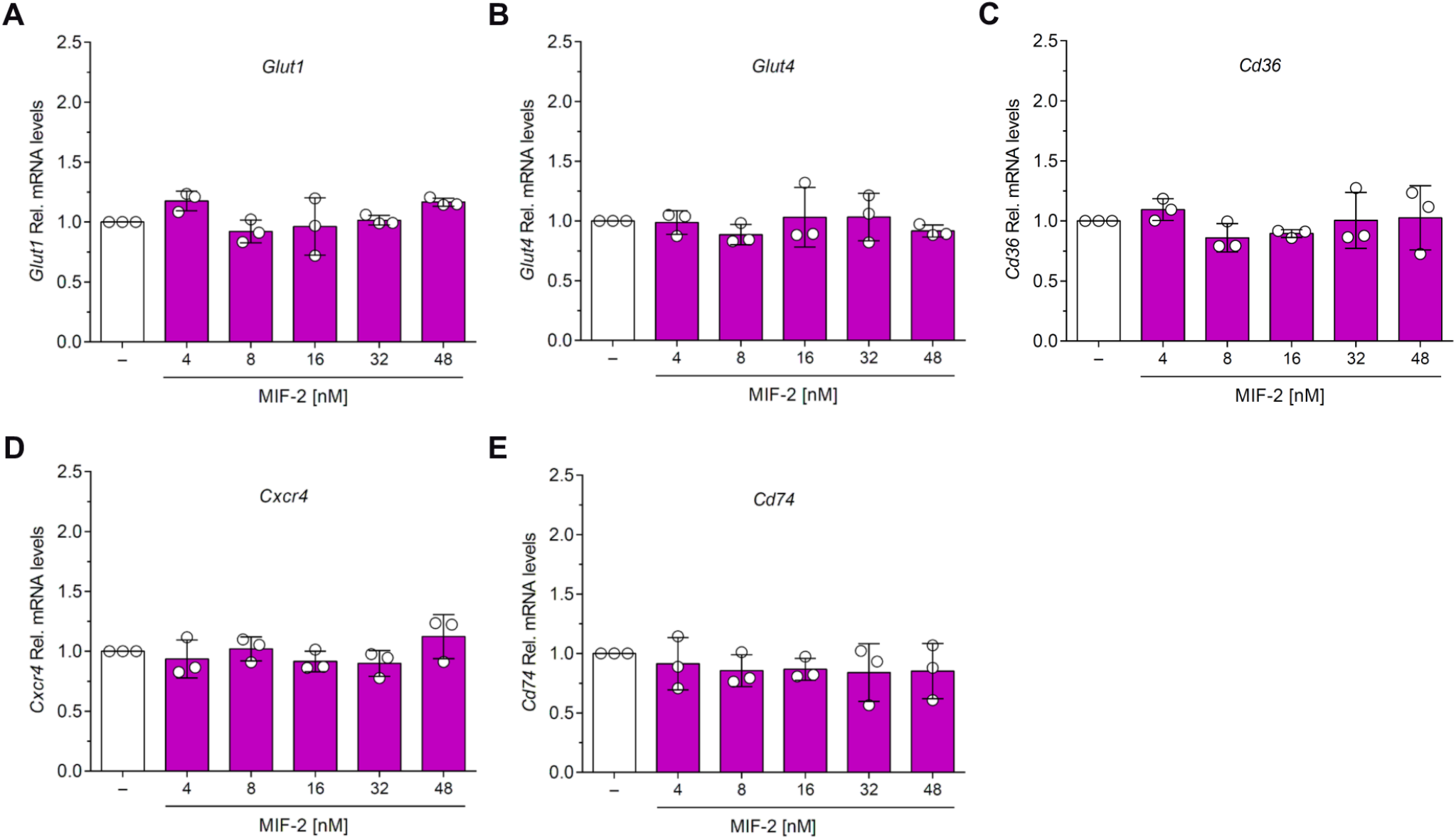
MIF-2 stimulation does not affect the expression of several lipid metabolism-related genes in Huh-7 cells. (A-C) Gene expression analysis of glucose transporters *Glut1 (A)* and *Glut4 (B) and* the scavenger receptor *Cd36 (C)* in Huh-7 cells after treatment with indicated concentrations of MIF-2. (D, E) Same as (A-C) except that the gene expression of the MIF/MIF-2 receptors *Cxcr4 (D)* and *Cd74 (E)* in Huh-7 cells was examined.

**Figure S10.**
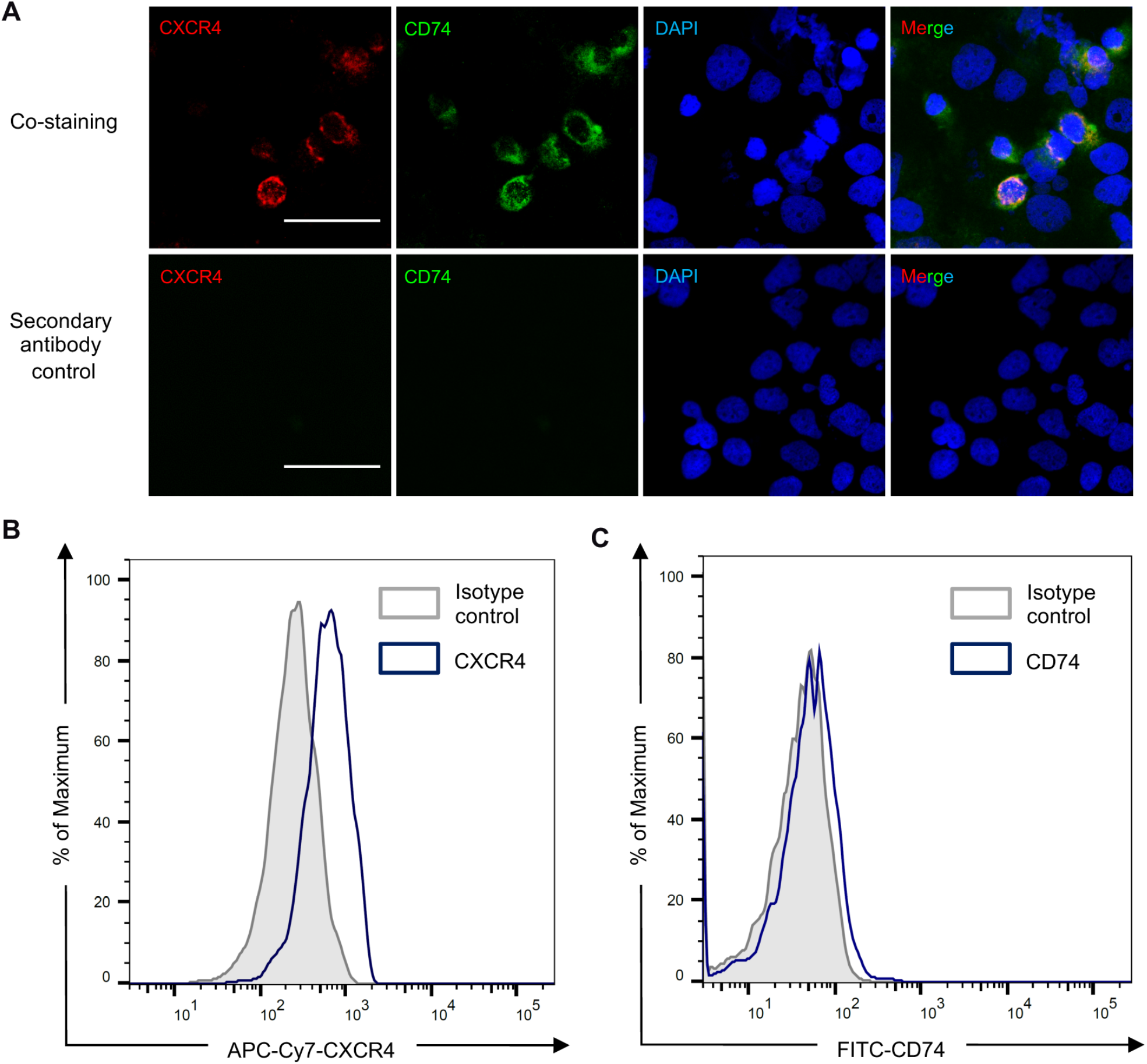
CXCR4 and CD74 are expressed in the human hepatocyte cell line Huh-7, as shown fluorescent staining and flow cytometry. (A) Colocalization of CXCR4 and CD74 in Huh-7 was visualized by immunostaining. Unstimulated cells were stained with anti-CXCR4 and anti-CD74 and corresponding fluorescently labeled secondary antibodies as indicated. Immunopositivity was detected by confocal microscopy. DAPI (blue) was used for counterstain. Scale bar, 50 µm. The secondary antibody control without primary antibody confirms specificity of the receptor expression signals. (B, C) Surface expression of CXCR4 and CD74 in Huh-7 as detected by flow cytometry. Unstimulated cells were stained with APC anti-CXCR4 (B) and FITC ani-CD74 (C), and expression was analyzed. Isotype control immunoglobulins served as negative control.

